# Distinct modulation of calcium-activated chloride channel TMEM16A by a novel drug-binding site

**DOI:** 10.1101/2023.08.06.552210

**Authors:** Jae Won Roh, Heon Yung Gee, Brian Wainger, Woo Kyung Kim, Wook Lee, Joo Hyun Nam

## Abstract

TMEM16A is a calcium-activated chloride channel with significant role in multiple cellular processes. Several TMEM16A inhibitors have been identified; however, their binding sites and inhibitory mechanisms remain unclear. Using magnolol and honokiol, the two regioisomeric inhibitors, as chemical probes, we have identified a novel drug-binding site distinct from the pore region, in TMEM16A, which is described here. With electrophysiology, unbiased molecular docking and clustering, molecular dynamics simulations, and experimental validation with mutant cycle analysis, we show that magnolol and honokiol utilize different drug-binding sites, pore and non-pore pockets. The pore blocker utilizes amino acids crucial for chloride passage, whereas the non-pore blocker allosterically modulates the pore residues to hinder ion permeation. Among 17 inhibitors tested, 11 were pore blockers and six were non-pore blockers, indicating the importance of this newly identified non-pore pocket. Our study provides insights into drug-binding mechanism in TMEM16A together with a rationale for future drug development.

## INTRODUCTION

Calcium-activated chloride channels (CaCCs) play important roles in physiological processes including epithelial fluid secretion, smooth muscle contraction, nociception transduction, cardiac activity control, capillary blood flow amplification, and diseases such as cancer progression, asthma, and hypertension^1–7^. Since the first successful cloning and expression of these channels including those in TMEM16 or anoctamin family^8–10^, many studies have been conducted on their electrophysiological and molecular features. Among members of the TMEM16 families, TMEM16A and TMEM16B are true pore-forming CaCCs^1, 11^.

Recent advances in the structural knowledge of TMEM16 proteins revealed that TMEM16A forms homodimers with 10 transmembrane (TM) helices and an ion-conducting pore surrounded by the TM3–TM8 helices within each subunit^12–14^. Upon increase in the cytosolic Ca^2+^ levels, two Ca^2+^ ions bind to the ‘principal’ Ca^2+^ binding interface of TM6–TM8, triggering conformation changes in TM6 and activating the ion-conducting state^13, 14^. TMEM16A activity is regulated by voltage, external Cl^−^, and internal Ca^2+^ ions^15^, it is also positively modulated by phosphatidylinositol 4,5-bisphosphate (PIP_2_) binding, and at least three sites on the cytoplasmic side are involved in channel gating and rundown^16–18^. In addition, a third Ca^2+^ ion binds between TM2 and TM10 and allosterically regulates channel opening^19^.

To date, many TMEM16A inhibitors have been identified, including T16inh-A01, CaCCinh-A01, Ani9, and 1PBC^2, 20–22^. While some drugs are specific to TMEM16A, others, such as benzbromarone or niclosamide^23^ are non-specific and capable of inhibiting other TMEM16 proteins such as TMEM16F^24^. Recent studies have utilized structure-based molecular docking and molecular dynamics (MD) simulation to show that the outer pore region is responsible for pore blockers such as CaCCinh-A01^25^ and 9-anthracene carboxylate (9-AC)^26^. Although these studies revealed the drug-binding site and interacting residues, important aspects of TMEM16A pharmacology still need to be addressed: (i) What are the roles of drug-interacting residues in normal channel function? (ii) Do multiple drug binding sites exist in TMEM16A? (iii) If so, what is the underlying molecular mechanism?

Searching for natural compounds that modulate TMEM16A activity, we identified magnolol and honokiol as TMEM16A inhibitors^27^. Interestingly, though magnolol and honokiol are structurally similar plant-derived regioisomers, honokiol is 4-fold more potent than magnolol. We probed whether this difference was due to the difference in binding affinity of these two molecules for TMEM16A, as suggested by the lock-and-key model^28^. Using electrophysiology, unbiased molecular docking and clustering, alanine scanning mutagenesis, and MD simulations, we showed that magnolol and honokiol bind to different drug-binding sites on TMEM16A, which we termed the pore pocket and the non-pore pocket, respectively. We also elucidated that the pore blocker molecule directly interacted with Cl^−^, while the non-pore blocker allosterically modulated the outer pore residues to inhibit TMEM16A activity. Finally, based on our analysis of the drug-binding sites, we classified the currently known TMEM16A inhibitors as pore or non-pore blockers. Overall, our work revealed new structural and pharmacological characteristics of the chemical regulation of TMEM16A.

## RESULTS

### Inhibition of TMEM16A currents by magnolol and honokiol

The structural difference between the regioisomeric compounds magnolol and honokiol is the position of attachment of one of the two hydroxyl group to one of the phenyl rings (Fig. 1a, f). The Tanimoto coefficient score for these two compounds is 95.2% (chemicals are generally considered similar when this value is >85%)^29^. We first verified the inhibitory effects of magnolol on TMEM16A Ca^2+^-activated Cl^−^ currents (Fig. 1b-d). In the whole-cell patch clamp configuration, the intracellular Ca^2+^ concentration was buffered at 300 nM, and magnolol was applied to the extracellular side. By utilizing step pulse protocol, we obtained similar half-maximal inhibitory concentrations (IC_50_) across tested voltage range, 28.71 ± 1.24 μM with Hill slope of 0.79 at 100 mV (Fig. 1e). We then conducted same experiments with honokiol, and again observed dose-dependent current inhibition (Fig. 1g-i). The IC_50_ value of honokiol increased only slightly at higher voltages, and was 6.60 ± 0.15 μM with Hill slope of 1.14 at 100 mV. Surprisingly, despite their similar structure, the inhibitory effects of magnolol and honokiol were substantially different, with honokiol being 4-fold more potent than magnolol.

**Fig. 1.**
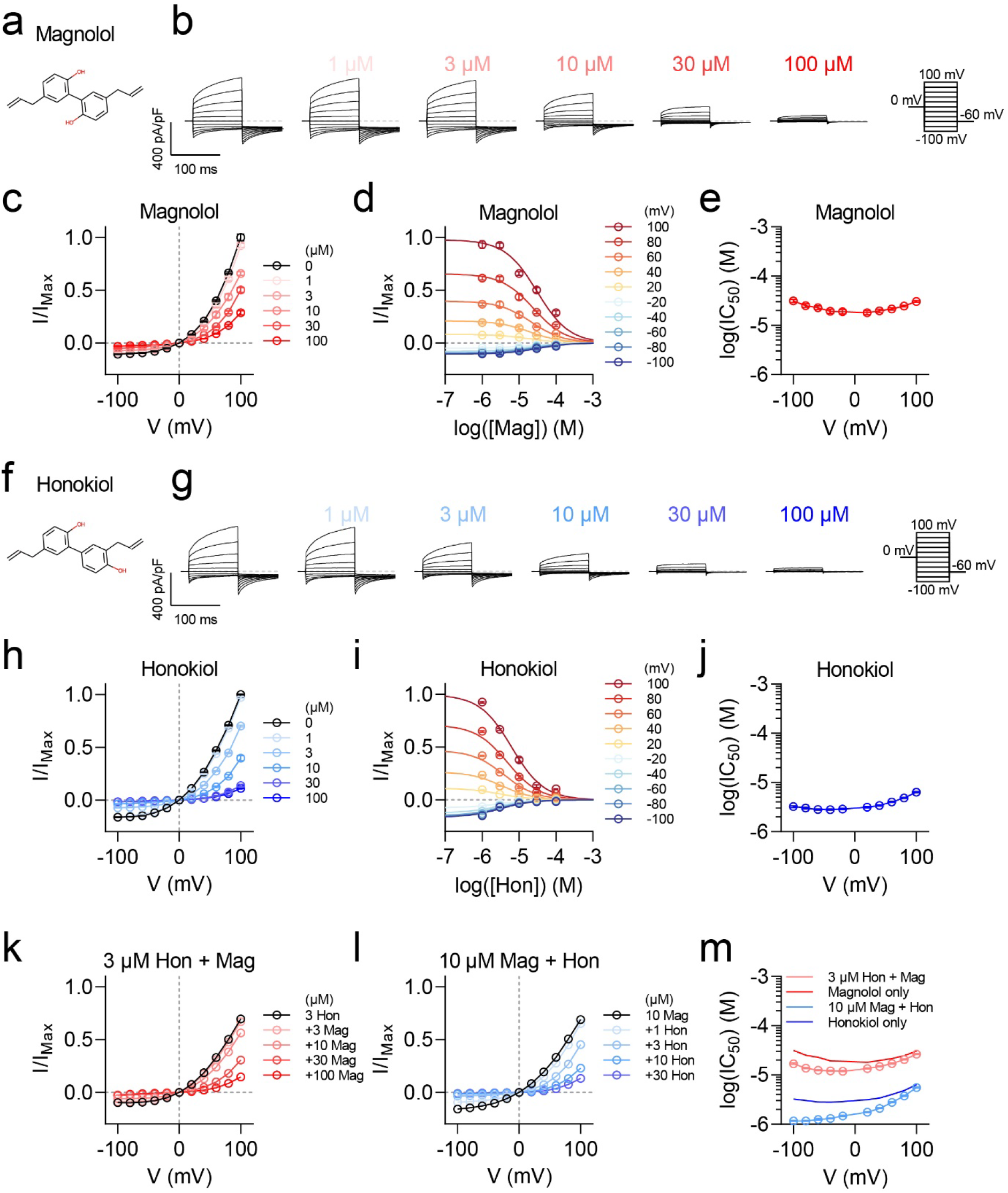
Inhibition of TMEM16A by magnolol and honokiol. **a.** 2D structure of magnolol (5,5’-diallyl-2,2’-biphenyldiol). **b.** Current traces of TMEM16A whole-cell currents upon magnolol administration on the extracellular side. Currents were evoked by 0 mV pre-pulse for 20 ms, then −100 to 100 mV test pulses for 100 ms, then −60 mV post-pulse for 100 ms. Intracellular Ca^2+^ was fixed at 300 nM. **c.** Current-voltage relationships of whole-cell currents upon magnolol administration. **d.** Dose-response relationships for magnolol in tested voltage range. The curves were obtained after fitting into the Hill equation. **e.** IC_50_ values calculated in **d** across tested voltage range. IC_50_ at 100 mV was 28.71 ± 1.24 μM with Hill slope of 0.79. **f.** 2D structure of honokiol (5,3’-diallyl-2,4’-dihydroxybiphenyl). **g.** Current traces of TMEM16A whole-cell currents upon honokiol administration on the extracellular side. Currents were evoked by 0 mV pre-pulse for 20 ms, then −100 to 100 mV test pulses for 100 ms, then −60 mV post-pulse for 100 ms. Intracellular Ca^2+^ was fixed at 300 nM. **h.** Current-voltage relationships of whole-cell currents upon honokiol administration. **i.** Dose-response relationships for magnolol in tested voltage range. The curves were obtained after fitting into the Hill equation. **j.** IC_50_ values calculated in **i** across tested voltage range. IC_50_ at 100 mV was 6.60 ± 0.15 μM with Hill slope of 1.14. **k.** Current-voltage relationships of whole-cell currents upon magnolol administration in the presence of 3 μM honokiol. **l.** Current-voltage relationships of whole-cell currents upon honokiol administration in the presence of 10 μM magnolol. **m.** IC_50_ values calculated in **k** and **l** across tested voltage range. *n* = 7 for magnolol administration in **c-e**; *n* = 7 for honokiol administration in **h-j**; *n* = 4 in **k**; *n* = 4 in **l**. Data are expressed as the mean ± SEM in **c**, **d**, **h**, **i**, **k**, **l**, and estimated value ± 95% confidence interval in **e**, **j**, **m**. Source data are provided as Source Data file.

We wondered whether magnolol and honokiol would compete with each other or synergistically work on blocking TMEM16A currents. Again, using the same whole-cell patch clamp configuration, the effect of magnolol was tested in the presence of 3 μM honokiol, and conversely, the effect of honokiol was tested in the presence of 10 μM magnolol (Fig. 1l, m). The concentrations of 3 and 10 μM were chosen for honokiol and magnolol, respectively because at these concentrations, the remaining current was about 70% of the maximum. In both conditions, the inhibitory effects of honokiol and magnolol increased at all tested voltage ranges, suggesting they do not competitively bind to TMEM16A but instead work in a synergistic manner (Fig. 1n).

### Identification of potential magnolol and honokiol-binding sites

Next, we aimed to identify the binding sites of magnolol and honokiol in TMEM16A. We conducted molecular docking using AutoDock Vina^28^ (henceforth referred to as Vina), one of the most widely used protein-ligand docking programs, and clustered the docking conformations using the *k*-means clustering algorithm (see Methods). To verify our molecular docking and clustering method, we used 1-hydroxy-3-(trifluoromethyl)pyrido[1,2-a]benzimidazole-4-carbonitrile (1PBC) and CaCCinh-A01, which are widely used TMEM16A inhibitors with known binding sites in the outer pore region (Supplementary Figs. 1 and 2). Indeed, our algorithm correctly identified the previously recognized binding sites^25, 30^. This gave us the confidence to use this algorithm to analyze the binding sites of magnolol and honokiol.

We applied our algorithm independently to magnolol and honokiol (Fig. 2a). Surprisingly, both magnolol and honokiol showed two similar clusters, one located on the extracellular end of the pore region and the other outside of the pore, henceforth named pore pocket and non-pore pocket according to their location (Fig. 2b). Interestingly, further analysis of each pocket by Vina revealed different binding characteristics. Magnolol bound more stably to the pore pocket, with binding energy (ΔG) of −7.0 kcal/mol as compared to the non-pore pocket (−6.0 kcal/mol, Fig. 2c). However, honokiol was more stable in the non-pore pocket, with a ΔG of - 7.5 kcal/mol compared to ΔG of −5.8 kcal/mol for the pore pocket (Fig. 2c). Interacting residues in the pore pocket were also significantly different between the two compounds; magnolol formed hydrogen bonds with R515 and E623, but not with E624 as did honokiol; whereas for honokiol, an interaction with E624 was observed with no interactions with R515 and E623 as for magnolol (Fig. 2c).

**Fig. 2.**
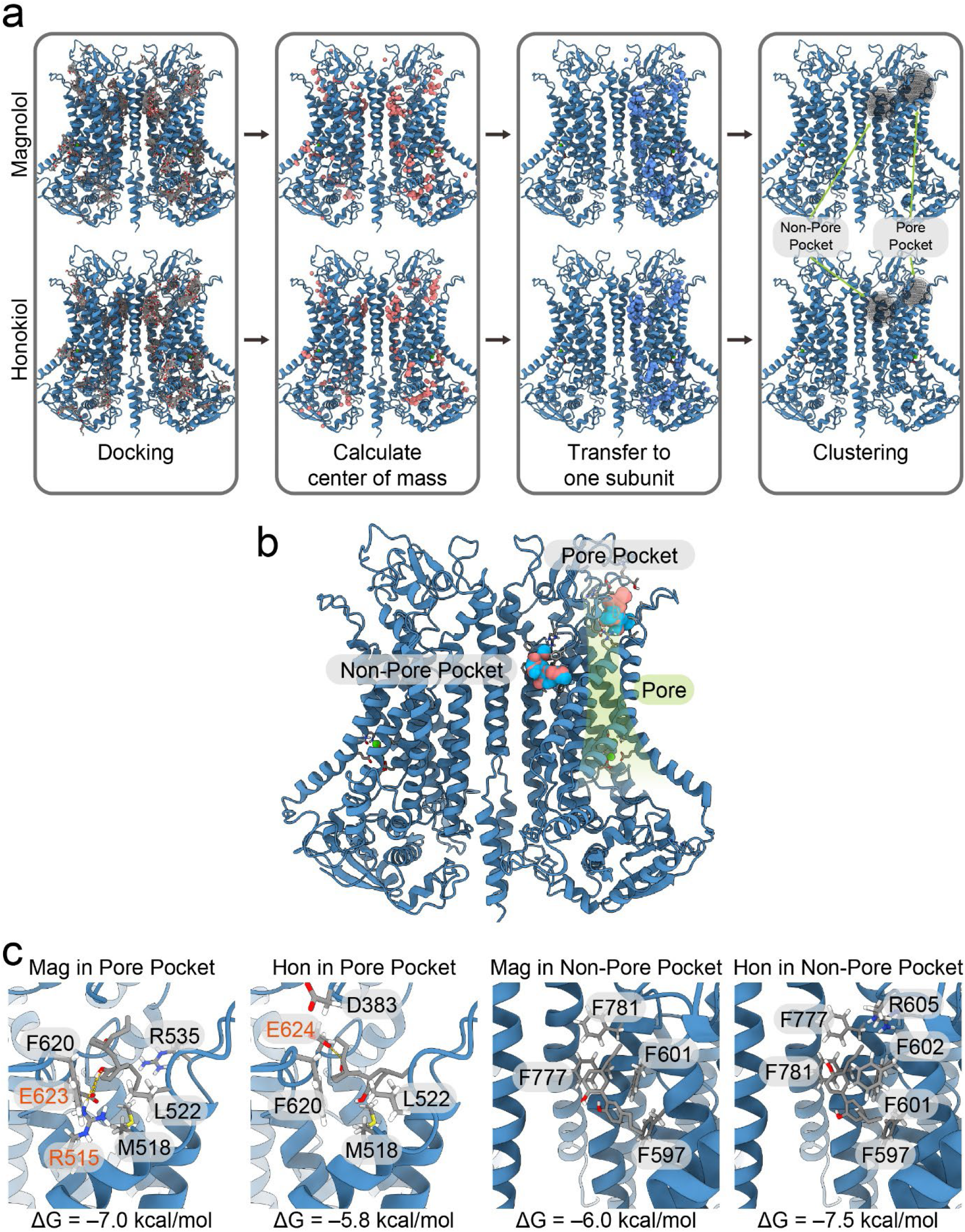
Identification of distinct magnolol and honokiol-binding sites. **a.** Molecular docking and clustering results for magnolol and honokiol. In the ‘clustering’ panel, each sphere represents ‘major’ clusters, and the size of the sphere is correlated with the number of docking poses. The detailed number of docking poses in each cluster is described in Supplementary Table 1. **b.** Space-filling atomic models of magnolol and honokiol binding in pore and non-pore pockets. Magnolol is colored red, honokiol is colored blue, and the pore region is shown in lime. **c.** Docking pose for magnolol and honokiol in the pore and non-pore pockets. The binding energy denoted as ΔG was calculated using AutoDock Vina^28^, and the interacting residues are shown as sticks, which were identified by PLIP (Protein-Ligand Interaction Profiler)^60^. Hydrogen bonds are shown as dotted yellow lines, and hydrogen bond-forming residue labels are colored orange.

### Alanine scanning mutagenesis in the pore and non-pore pockets

We examined whether the different binding affinities and binding sites of magnolol and honokiol could also be distinguished by patch clamp electrophysiology. For this we examined the involvement of all the relevant residues by alanine (Ala; A) scanning mutagenesis by mutating each residue to Ala, since alanine has a small side chain and can maintain stable secondary structures in proteins, making it ideal for this type of studies. A reduced binding affinity of an Ala mutant would indicate involvement of that particular residue in the compound-binding, which in turn should indicate a decreased inhibition by the test compound. Based on the previous knowledge of the binding characteristics (Fig. 2c), we selected the following residues: D383, R515, M518, L522, R535, K603, F620, E623, E624, and E633 from the pore pocket; and F597, F601, F602, R605, F777, and F781 from the non-pore pocket (Fig. 3a, b).

**Fig. 3.**
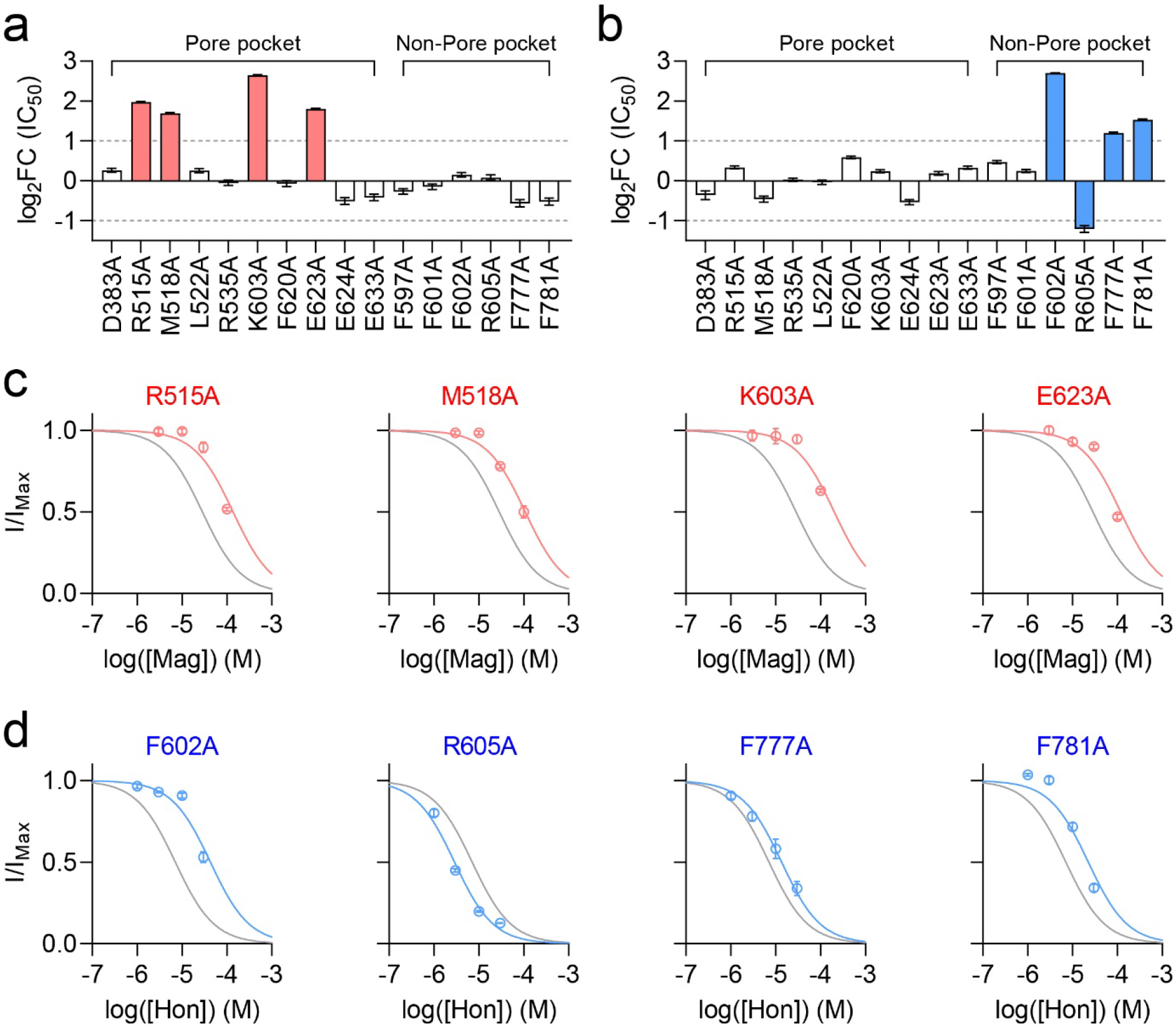
Alanine-scanning mutagenesis in the pore and non-pore pockets. **a.** Alanine scanning mutagenesis results for magnolol. **b.** Alanine scanning mutagenesis results for honokiol. In **a** and **b**, intracellular Ca^2+^ was fixed at 300 nM in a whole-cell patch clamp configuration, and magnolol or honokiol was administered in the extracellular side. For each mutant, IC_50_ at 100 mV was calculated by fitting to the Hill equation, and fold-change was calculated by dividing by IC_50_ of the WT. The gray lines indicate a 2-fold change. Mutants with more than two-fold-change in IC_50_ are colored in either red or blue. *n* = 4 for each mutant in **a** and **b**. Current-voltage relationships, dose-response relationships, and IC_50_ for each voltage are shown in Supplementary Figs. 3-8. **c.** Dose-response relationships for R515A, M518A, K603A, and E623A mutants upon magnolol administration at 100 mV. Gray line indicates dose-response relationship for the WT. d. Dose-response relationships for F602A, R605A, F777A, and F781A mutants upon honokiol administration at 100 mV. Gray line indicates dose-response relationship for the WT. Current-voltage relationships, dose-response relationships, and IC_50_ for each voltage are shown in Supplementary Figs. 10-15. Data are expressed as the estimated value ± 95% confidence interval in **a**, **b** and mean ± SEM in **c**, **d**. Source data are provided as Source Data file.

Each Ala mutant was treated with different concentration of either magnolol or honokiol on the extracellular side in the whole-cell patch clamp. IC_50_ values calculated by fitting the Hill equation at 100 mV was divided by IC_50_ of the wild type (WT) to calculate fold-change of each mutant. Magnolol showed significantly decreased inhibition rate in R515A, M518A, K603A, and E623A mutants, with IC_50_ value of 120.4 ± 3.6 μM, 99.0 ± 2.0 μM, 191.9 ± 9.1 μM, and 110 ± 3.2 μM, respectively (Fig. 3a, c, Supplementary Figs. 3-5). Incidentally, all these residues were located in the pore pocket. On the other hand, honokiol showed significantly decreased inhibition rate for F602A, F777A, and F781A mutants, with IC_50_ value of 41.2 ± 1.4 μM, 14.6 ± 0.6 μM, and 18.4 ± 0.5 μM, respectively. Also, for R605A mutant, IC_50_ value was lower than WT with 2.76 ± 0.05 μM (Fig. 3b, d, Supplementary Figs. 6-8). All the analyzed residues were part of the non-pore pocket. Overall, our mutagenesis experiments demonstrated that magnolol and honokiol act on different pockets, which is consistent with our pocket-specific docking results. In addition, magnolol and honokiol are now referred to as the pore blocker and non-pore blocker, respectively, as experimentally determined.

### Side chain contribution in the pore and non-pore pockets

We carried out further experiments to characterize the residues that significantly shifted IC_50_ compared to the WT (Fig. 3c, d). Four residues were chosen from each pocket, namely R515, M518, K603, and E623 in the pore pocket, and F602, R605, F777, and F781 in the non-pore pocket. We conducted additional mutagenesis and patch clamp experiments to examine the impact of side chains. Arg in the positions 515 and 605 were mutated as follows: to Lys for same positive charge but with smaller side chain, to Ile for a hydrophobic side chain larger than Alanine, and to Asp for the converting the positive charged side chain to a negatively charged one. Met518 was mutated to Thr, Val, and Ile, as each of these residue are only slightly different with more or less bulky hydrophobic side chains. Lys603 was mutated to Arg, Ile, and Asp, similar to R515. Glu623 was mutated as follows: Asp for same charge but with smaller side chain size, Ile for hydrophobic side chain, and Lys for charge reversal. Three Phe of the four residues in the non-pore pocket, F602, F777, and F781 were mutated as follows: Tyr for more hydrophilic side chain with similar size, Ile for hydrophobic side chain, and Trp for more bulky side chain. To summarize, all of the amino acid residues that showed significant differences from the wild type TMEM16A when mutated to Ala were further tested with three additional side chains with different properties.

Among the R515 mutants, substitution with Lys (R515K) showed similar dose-response curve as in the WT, while R515I and R515D mutants were similar to R515A (Fig. 4a). In case of M518 mutants, M518T was similar to the dose-response curves of the WT, and M518I and M518V mutants showed decreased response to magnolol, similar to M518A (Fig. 4a). It is worthwhile to note that in mTMEM16A, Met in 518 position is substituted by Thr. In K603 position, all of additional mutants, K603R, K603I, and K603D, were significantly different from the WT (Fig. 4a). Finally, in the E623 position, the dose-response curve of E623D mutant was located between the WT and E623A, while E623I and E623K mutants were similar to E623A (Fig, 4a, Supplementary Figs. 9-11).

**Fig. 4.**
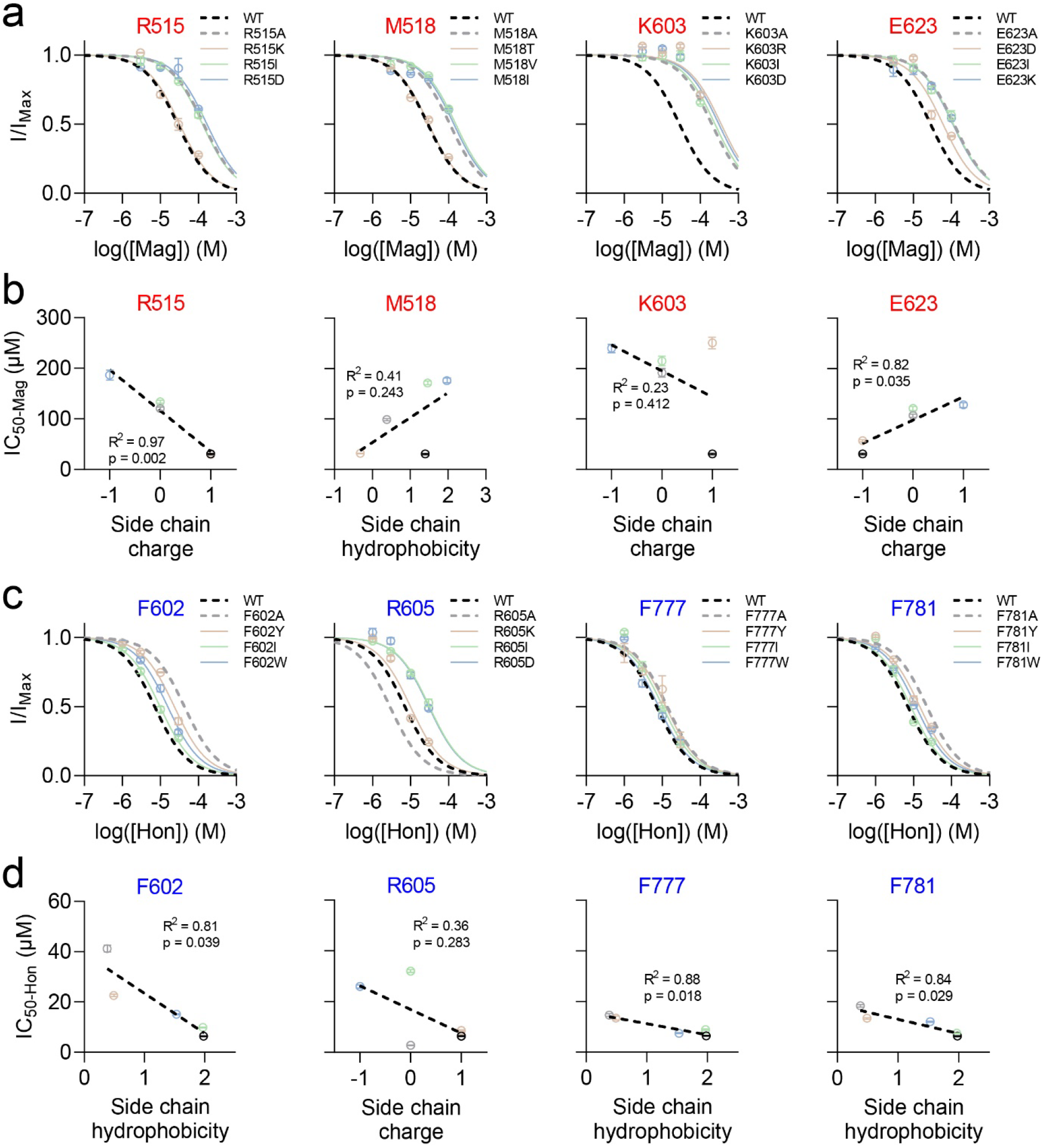
Side chain contribution of the pore and non-pore pocket residues. **a.** Dose-response relationships for magnolol with various mutants in R515, M518, K603, and E623 positions. Intracellular Ca^2+^ was fixed at 300 nM in a whole-cell patch clamp configuration, and magnolol was administered in the extracellular side. The curves are fit to the Hill equation. **b.** Relationship of magnolol IC_50_ with either side chain charge or side chain hydrophobicity. **c.** Dose-response relationships for honokiol with various mutants in F602, R605, F777, and F781 positions. Intracellular Ca^2+^ was fixed at 300 nM in a whole-cell patch clamp configuration, and honokiol was administered in the extracellular side. In **b** and **d**, Side chain hydrophobicity value was adopted from Zhao & Erwin^31^. In **a** and **c**, *n* = 4 for each mutant. Data are expressed as the mean ± SEM in **a**, **c** and estimated value ± 95% confidence interval in **b**, **d**.

We then plotted IC_50_ values of various mutants and the WT TMEM16A against either the side chain charge or side chain hydrophobicity, depending on the original residue type: if original side chain had positive or negative charge, then side chain charge was plotted, and if original side chain was neutral, then side chain hydrophobicity was plotted. The side chain hydrophobicity was adopted from Zhao & Erwin^31^. The effect of magnolol inhibition on R515 and E623 showed significant linear relationship, indicating side chain charges of these two residues was major factor in magnolol-binding (Fig. 4b). M518 showed no significant relationship with change in type of side chain hydrophobicity (Fig. 4b). In case of K603, none of the mutations we tested were similar to the WT, indicating that both side chain charge and size are important (Fig. 4b).

We adopted same strategy for the non-pore pocket mutants. Mutations in F602, F777, and F781 positions showed various degree of honokiol inhibition, while in R605 position, mutations other than R605A hindered honokiol inhibition (Fig. 4c, Supplementary Figs. 12-14). Plotting IC_50_ with side chain hydrophobicity in F602, F777, and F781 showed strong linear relationship, indicating that hydrophobic interaction was major type of molecular interaction (Fig. 4d). For the residue R605, there was no clear relationship between side chain charge and IC_50_ (Fig. 4d). In summary, our additional experiments on side chain mutagenesis indicated that in the pore pocket, the side chain charge plays a crucial role, whereas in the non-pore pocket, the side chain hydrophobicity is the primary factor.

### Closed TMEM16A pore upon drug binding

To gain molecular insights into the inhibition mechanisms of pore and non-pore blockers, we performed MD simulations. The Ca^2+^-bound TMEM16A structure was used as a control, and pore and non-pore blockers were docked into each pocket prior to the simulation using Vina. Each condition was run in triplicate, and root mean square deviation (RMSD) of TMEM16A and the drugs (either magnolol or honokiol) were stable throughout MD simulations (Fig. 5a, b). The contact probability and the decomposition of energy showed similar results with our mutant electrophysiology data (Supplementary Fig. 15); the residues R515, M518, and E623 of TMEM16A had a contact probability of >0.5 with the pore blocker, and negative (favorable) binding energy; F602, F777, and F781 showed >0.5 contact probability with the non-pore blocker and a negative binding energy, while R605 showed a high contact probability but positive (unfavorable) binding energy with the non-pore blocker, partially explaining the increased non-pore blocker effect in the R605A mutant (Fig. 3b).

**Fig. 5.**
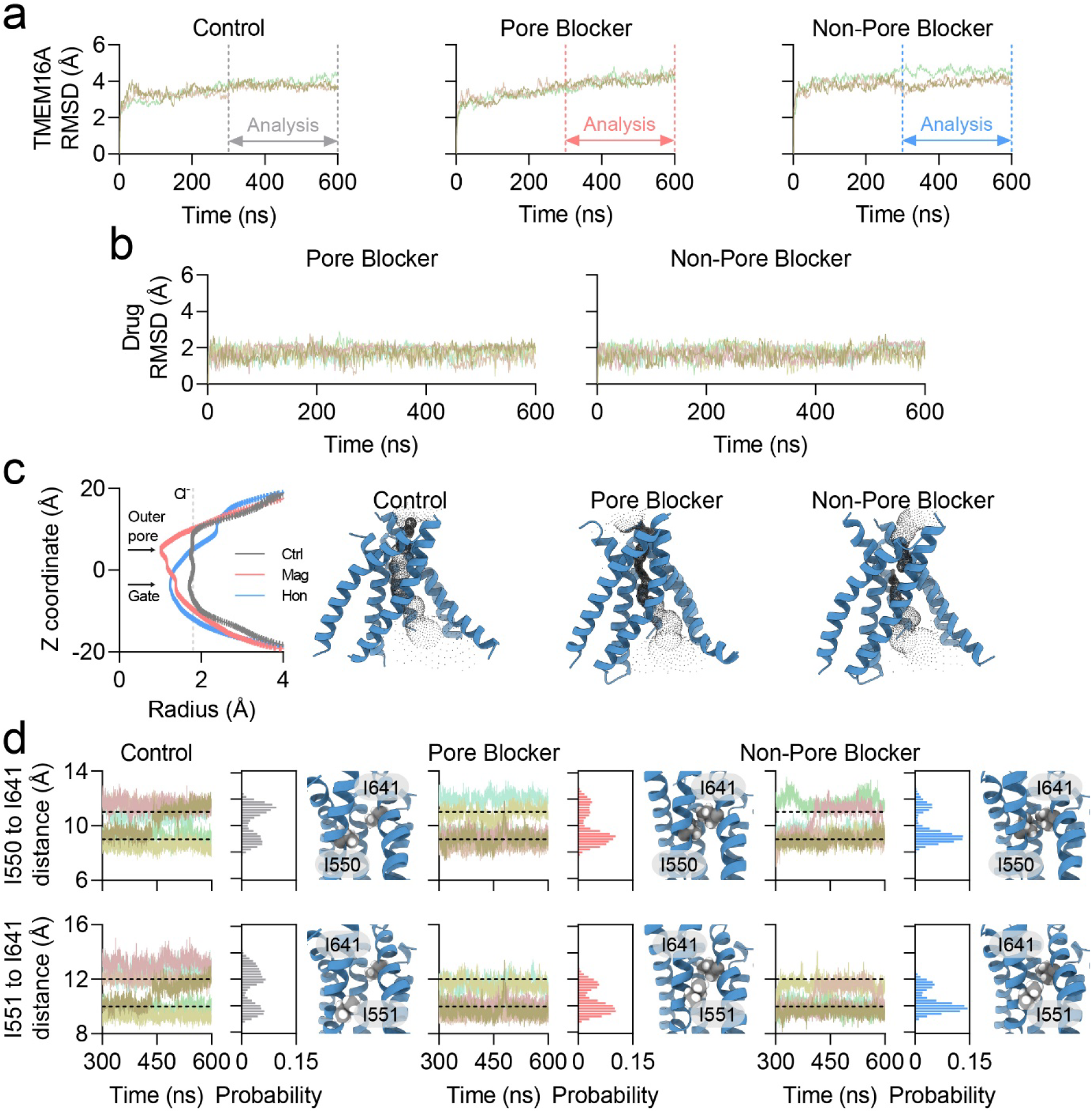
Collapsed TMEM16A channel pore upon drug binding. **a.** TMEM16A root mean square deviation (RMSD) calculation during MD simulations. Each simulation was run in triplicate. 300 to 600 ns were used for further analysis, which is indicated. **b.** Drug RMSD calculation during MD simulations. **c.** Ion-permeating pore region identified by the HOLE^32^ in MD simulations. The locations of the outer pore region and gate are indicated by the arrows. The gate refers to the I550-I641 and I551-I641 interaction sites. The dotted gray line indicates the radius of Cl^−^ ion. **d.** Time course, frequency distribution, and the molecular model of the pore structure from each simulation condition. The snapshot that was most similar to **c** was selected using the least-squares method. Frequency distribution of the I550-I641 or I551-I641 gate was obtained by calculating the center of mass of the I550, I551 and I641 Cα atoms, and the bin size was 0.2 Å. Data are expressed as the mean ± SEM. Source data are provided as Source Data file.

We also calculated the channel pores using HOLE^32^ and observed that TM helices 3 to 8 formed a water-accessible pore region (Fig. 5c). The pore of ‘control’ Ca^2+^-bound TMEM16A structure showed comparable radius to Cl^−^ ion. Surprisingly, both pore and non-pore blocker-bound structures showed collapsed channel pores with radii smaller than those of Cl^−^ ions (Fig. 5c). We then examined the steric gates in the middle of the pore, which were recently identified as two pairs of residues, I550-I641 and I551-I641^33^ (Fig. 5d). Both gates were significantly closed by each blocker. The most frequently observed gating distances in the drug-free state were 11.4 Å for I550-I641 and 12.0 Å for I551-I641 (Fig. 5d). When the pore blocker was bound, the gating distances were 9.2 Å and 9.6 Å, respectively, and in presence of the non-pore blocker, the distances were 9.2 Å and 9.8 Å respectively (Fig. 5d). Unlike differences in the gating distances, the difference in the angle of TM6 was observed to be the major difference between Ca^2+^-free and Ca^2+^-bound structures in earlier studies^14^, which was not found to be significantly different in the MD simulations in the present study (Supplementary Fig. 16).

During the MD simulations, structural rearrangements were observed in the outer pore region (Supplementary Fig. 17). When the pore blocker was bound, TM3 was closer to TM4 by 2.2 Å, and TM4 was closer to TM6 by 6.0 Å. These changes in TM3 and TM4 were responsible for the major constriction in the outer pore region caused by the direct binding of magnolol in the pore pocket. In contrast, when the non-pore blocker was bound, TM5 was closer to the center of the pore by 1.6 Å, and TM4 shifted away from the pore by 3.3 Å. These changes were presumably observed because the direct-binding site of the non-pore blocker is in TM5 (Supplementary Fig. 17).

We performed additional MD simulations where pore-blocker was bound to the R515A mutant and non-pore blocker was bound to the F602A/F777A/F781A triple mutant (Supplementary Fig. 18). In all of the simulations runs, drugs (either pore or non-pore blocker) dissociated from TMEM16A in less than 300 ns, again indicating the importance of these residues in the respective pockets (Supplementary Fig. 18).

### Electrostatic interactions in the pore pocket

The closure of I550-I641 and I551-I641 gates seemed responsible for the inhibitory effects of the pore and non-pore blockers. We decided to further investigate the role of the outer pore residues based on the following observations: (i) the outer pore structural arrangement was significantly different in presence of pore and non-pore blocker (Fig. 5c, Supplementary Fig. 17); (ii) mutagenesis of residues R515, K603, E623, and E633 of the outer pore region to Ala showed impaired channel function (Supplementary Fig. 19). The impaired channel function was assessed by decreased whole-cell current density, weak time-dependent activation on high depolarizing voltages, and increased tail current deactivation (Supplementary Fig. 19). Interestingly, these residues were also responsible for binding of the pore blocker (Fig. 2-4). To identify the function of these residues in the normal channel function, we fitted normalized IV curves obtained by saturating Ca^2+^ in the ion permeation equation developed by the Dutzler group^34^ (Fig. 6a-c; also see Methods). When fitted to the three-barrier model, the WT TMEM16A showed high extracellular and cytoplasmic energy barriers and a lower middle barrier, similar to previously reported results^35^ (Fig. 6d). We then performed the same analysis on the R515A, K603A, E623A, and E633A mutants (Fig. 6e). R515A and K603A mutants exhibited a higher middle barrier than the WT, implying direct involvement in Cl^−^ permeation. Surprisingly, the E623A and E633A mutants showed outwardly rectifying IV curves, which were calculated to have a higher middle energy barrier. This was counterintuitive because removing negative charges in E623A and E633A mutants should facilitate the movement of negatively charged Cl^−^ ions along the pore. The increase in the energy barrier of the E623A and E633A mutants led us to hypothesize that these residues may be involved in modulation of other positively charged amino acids. Indeed, E623 and E633 are located very close to R515 and/or K603.

**Fig. 6.**
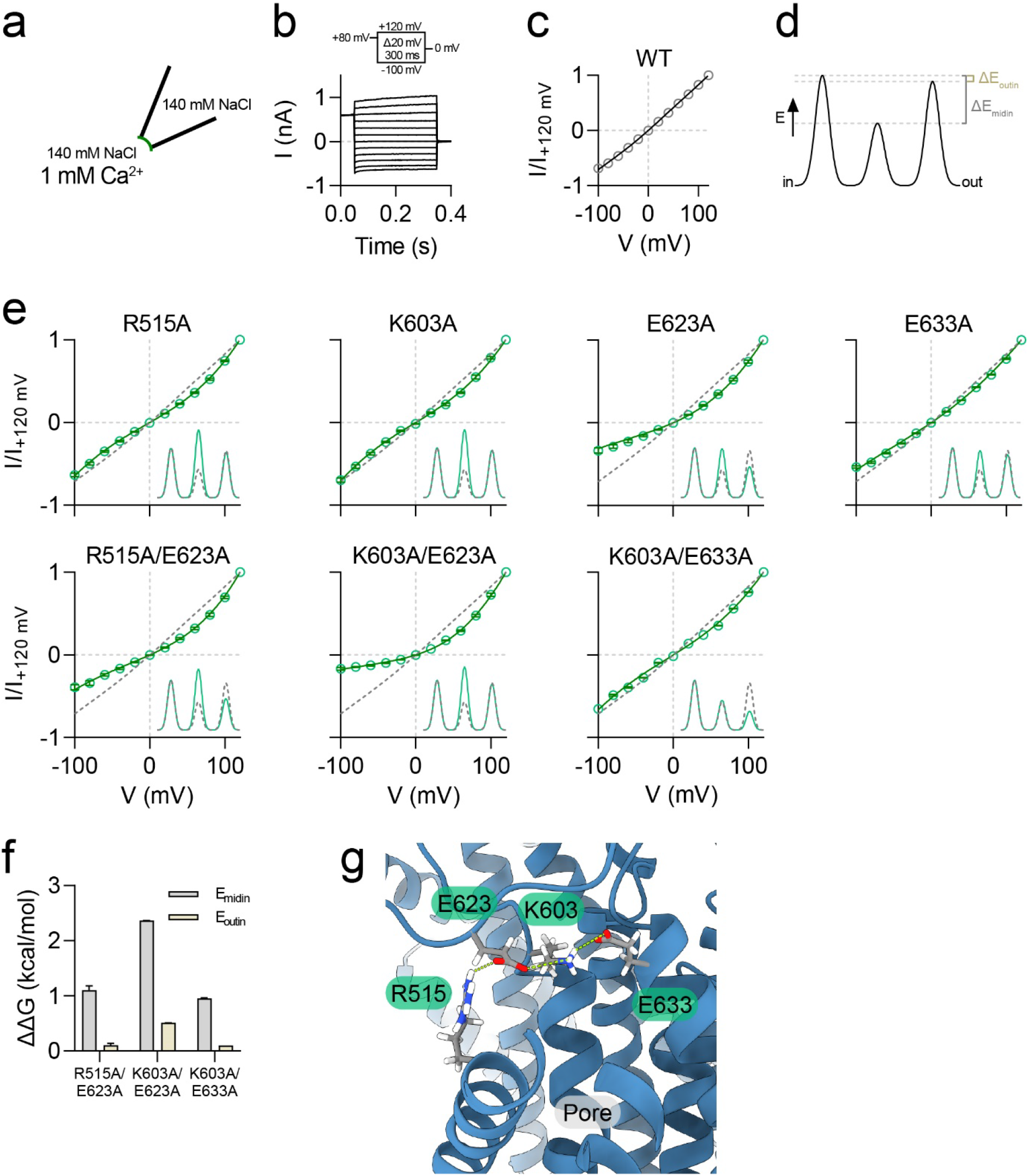
Electrostatic interactions in the pore pocket for Cl^−^ permeation and drug-binding. **a.** Illustration of the inside-out patch clamp configuration for the ion permeation model calculations. 1 mM Ca^2+^ was used to activate TMEM16A in symmetrical 140 mM NaCl solution in the inside-out patch clamp configuration. **b.** Current trace of WT TMEM16A activation curve. Currents were evoked by 80 mV pre-pulse for 50 ms, then –100 to 120 mV test pulses for 300 ms. **c.** Normalized current trace for WT TMEM16A. Gray circles represent the normalized current, and the black line is the fit to the ion permeation model (see Methods). *n* = 22. **d.** Energy barrier of WT TMEM16A. The energy barriers were calculated by fitting the result from **c** to the ion permeation model (see Methods). ΔE_midin_ is –0.8678 ± 0.0032 kcal/mol and ΔE_outin_ is –0.1104 ± 0.0002 kcal/mol. **e.** Normalized current traces for single and double mutants. Green circles represent the normalized current of each mutant, and the green line is the fit to the ion permeation model. The dotted gray line is for the WT. An energy barrier was observed at the inlet. *n* = 18 for R515A; *n* = 17 for E623A; *n* = 10 for E633A; *n* = 9 for R515A/E623A; *n* = 8 for K603A/E623A; *n* = 10 for K603A/E633A. **f.** ΔΔG values calculated from double-mutant cycle analysis (see Supplementary Fig. 27 and Methods). The E_midin_ values for R515A/E623A, K603A/E623A, and K603A/E633A were 1.1050 ± 0.1308, 2.3649 ± 0.0045, and 0.9563 ± 0.0089 kcal/mol, respectively. The E_outin_ values for R515A/E623A, K603A/E623A, and K603A/E633A are 0.1018 ± 0.0618, 0.5088 ± 0.0008, and 0.0994 ± 0.0018 kcal/mol, respectively. **g.** Locations of R515, K603, E623, and E633 in the outer pore region. Negatively charged oxygen atoms are shown in red, positively charged nitrogen atoms are shown in blue, and potential electrostatic interactions are shown in yellow. Data are expressed as mean ± SEM in **c** and **e** and estimated value ± 95% confidence interval in **f**. Source data are provided as Source Data file.

To directly measure their electrostatic interactions, we calculated the energy barriers of the double mutants, R515A/E623A, K603A/E623A, and K603A/E633A, and performed double-mutant cycle analysis by fitting IV curves (Fig. 6e, f; also see Methods). The double-mutant cycle analysis allowed us to calculate the electrostatic energy coupling of two residues; when the ΔΔG value deviated from 0, the two residues were likely to interact. Interestingly, R515-E623, K603-E623, and K603-E633 were found to be energetically coupled, with ΔΔG value of 1.10 ± 0.13 kcal/mol, 2.36 ± 0.00 kcal/mol, 0.96 ± 0.01 kcal/mol at the middle barrier, respectively, and 0.10 ± 0.06 kcal/mol, 0.51 ± 0.00 kcal/mol, 0.10 ± 0.00 kcal/mol at the outer barrier. Unlike pore residues, the non-pore pocket residues, F602 and F777, which are important for non-pore blocker-binding, were energetically independent in both E_midin_ and E_outin_ (Supplementary Fig. 20).

Overall, our results suggest that R515, K603, E623, and E633 are important for drug binding and Cl^−^ permeation in the pore pocket, which explains the inhibition mechanism of pore pocket blockers in the outer pore region; direct interactions with R515 and K603 with the drug, block Cl^−^ interactions with these residues, and E623 and E633 interact with R515 and/or K603, presumably help in stabilizing R515 and K603 for Cl^−^ permeation (Fig. 6g).

### Broken salt bridges in the presence of non-pore blocker

We then investigated whether previously observed structural changes in the outer pore region by non-pore blocker-binding were related to R515-E623-K603-E633 interaction differences. Thus, we analyzed the salt bridges formed by R515/E623, K603/E623, and K603/E633 in our MD simulation trajectories (Fig. 7a). The probability of R515/E623 salt bridge formation was lower when the pore-blocker was bound, but higher in the presence of the non-pore blocker (Fig. 7b). The K603/E633 bridge was increased in the pore blocker and almost same when the non-pore blocker was bound (Fig. 7b). The K603/E633 bridge was decreased in both pore and non-pore blockers (Fig. 7b).

**Fig. 7.**
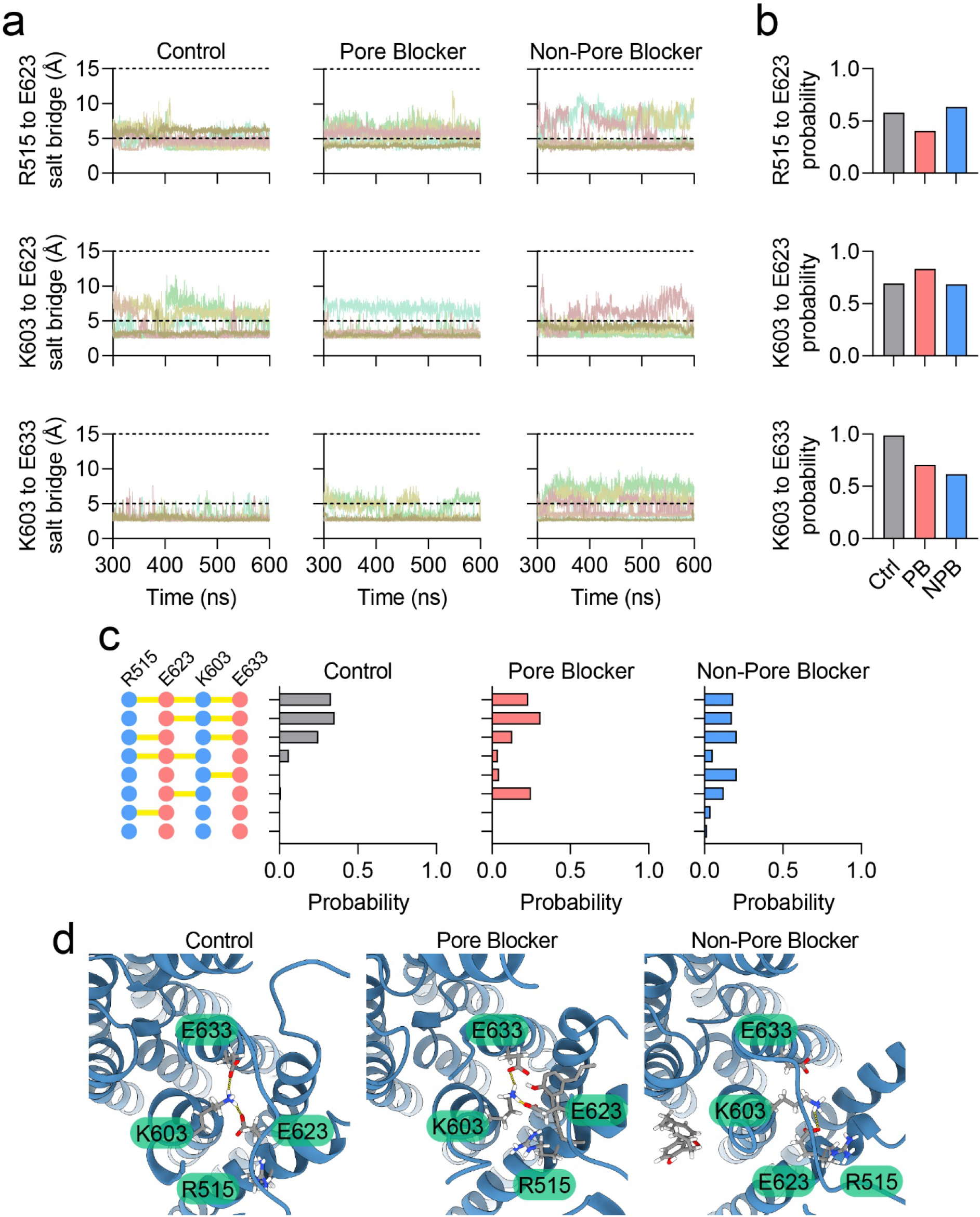
Disruption of the electrostatic interactions in the pore pocket by non-pore blocker. **a.** Time course of R515-E623, K603-E623, and K603-E633 salt bridge distance in MD simulations. Salt bridges were calculated in VMD^58^ and the cutoff distance was 5 Å. When there was no salt bridge formation, the distance was designated as 15 Å for visual purpose. **b.** Probability of R515-E623, E623-K603, and K603-E633 salt bridge formation in MD simulations. **c.** Probability of R515-E623, E623-K603, and K603-E633 salt bridge configurations in MD simulations. Source data are provided as Source Data file.

We next aimed to analyze salt bridge conformations in each MD simulation condition. There were 8 (2^3^) different interaction possibilities for each salt bridge formed, or not formed. For simple annotation, we designated a formed bridge as ‘1’ and an unformed bridge as ‘0’ and listed them in the order, R515/E623–E623/K603–K603/E633; for example, for unformed R515/E623, formed E623/K603, and formed K603/E633 bridges, the annotation would be ‘0–1–1’ (Fig. 7c). For drug-free control TMEM16A trajectories, 0–1–1 conformation was the most common (35.1%), followed by 1-1-1 conformation (32.8%) (Fig. 7c). The pore blocker-bound TMEM16A also showed 0–1–1 as the most common conformation (30.9%), followed by 0–1–0 (24.7%) (Fig. 7c). Strikingly, non-pore blocker-bound TMEM16A was largely different from the others; 0–0–1 was the most common (20.6%), followed by 1–0–1 (20.5%), 1–1–1 (18.4%), 0–1–1 (17.4%) (Fig. 7c). The K603/E633 bridge was impaired in the honokiol-bound TMEM16A structure compared to that in the control group, and both K603 and R515 were dislocated from the original position (Fig. 7d). Indeed, when we treated protein with 10 μM non-pore blocker and fitted the ion permeation equation of TMEM16A, the middle energy barrier increased, while the outer barrier remained the same as that of the WT, thus partially resembling R515A and K603A mutants (Supplementary Fig. 21).

With these experiments, we demonstrated the molecular mechanism of the non-pore blocker on the outer pore region; disruption of R515-E623-K603-E633 interactions in the pore pocket causes dislocation of K603 and R515, which may hinder Cl^−^ permeation.

### Classification of TMEM16A inhibitors

To date, many TMEM16A inhibitors have been identified through large library screening^2, 20–23^, but their binding sites are mostly unknown, except for CaCCinh-A01^25^, 1PBC^30^, and 9-AC^33^. All three drugs bind to the outer pore region, which is identical to the pore pocket in our study. We wondered whether other TMEM16A inhibitors might utilize the non-pore pocket identified in this study. We selected widely used TMEM16A-specific inhibitors for classification by pocket specificity, such as T16inh-A01, CaCCinh-A01, Ani9, and 1PBC^20–22^; classical anion channel blockers, 4,4’-diisothiocyanatostilbene-2,2’-disulfonic acid (DIDS), niflumic acid, 5-nitro-2-(3-phenylpropylamino)benzoic acid (NPPB), and N-phenylanthracilic acid (NPA)^36, 37^; inhibitors identified by screening FDA-approved drugs, dichlorophen, hexachlorophene, benzbromarone, and niclosamide^2, 23^; and other CaCC blockers such as mefloquine, fluoxetine, and tamoxifen^9^. We selected R515A as the mutant for the pore pocket because it was implicated in the binding of magnolol as well as 1PBC^30^ and CaCCinh-A01^25^. As the non-pore mutant, we chose F602A/F777A double mutant because this mutant decreased the effect of honokiol to 5-fold (Supplementary Fig. 22). Each drug was tested at 10 μM concentration on the WT, pore mutant (R515A), and non-pore mutant (F602A/F777A) TMEM16A channels (Fig. 8). The difference in the inhibition rate of the WT compared to each mutant was calculated to measure the fraction of inhibitory effects on the pore and non-pore pockets. We plotted the inhibitory rate of each drug with the calculated fraction of the pocket, which allowed us to visually observe the preference of each drug on the pore or non-pore pocket (Fig. 8). Interestingly, magnolol, T16inh-A01, CaCCinh-A01, Ani9, 1PBC, DIDS, NPA, NPPB, niflumic acid, mefloquine, and tamoxifen favored the pore pocket, whereas honokiol, dichlorophen, hexachlorophene, benzbromarone, and niclosamide more favorably bound to the non-pore pocket (Fig. 8, Supplementary Figs. 23 and 24).

**Fig. 8.**
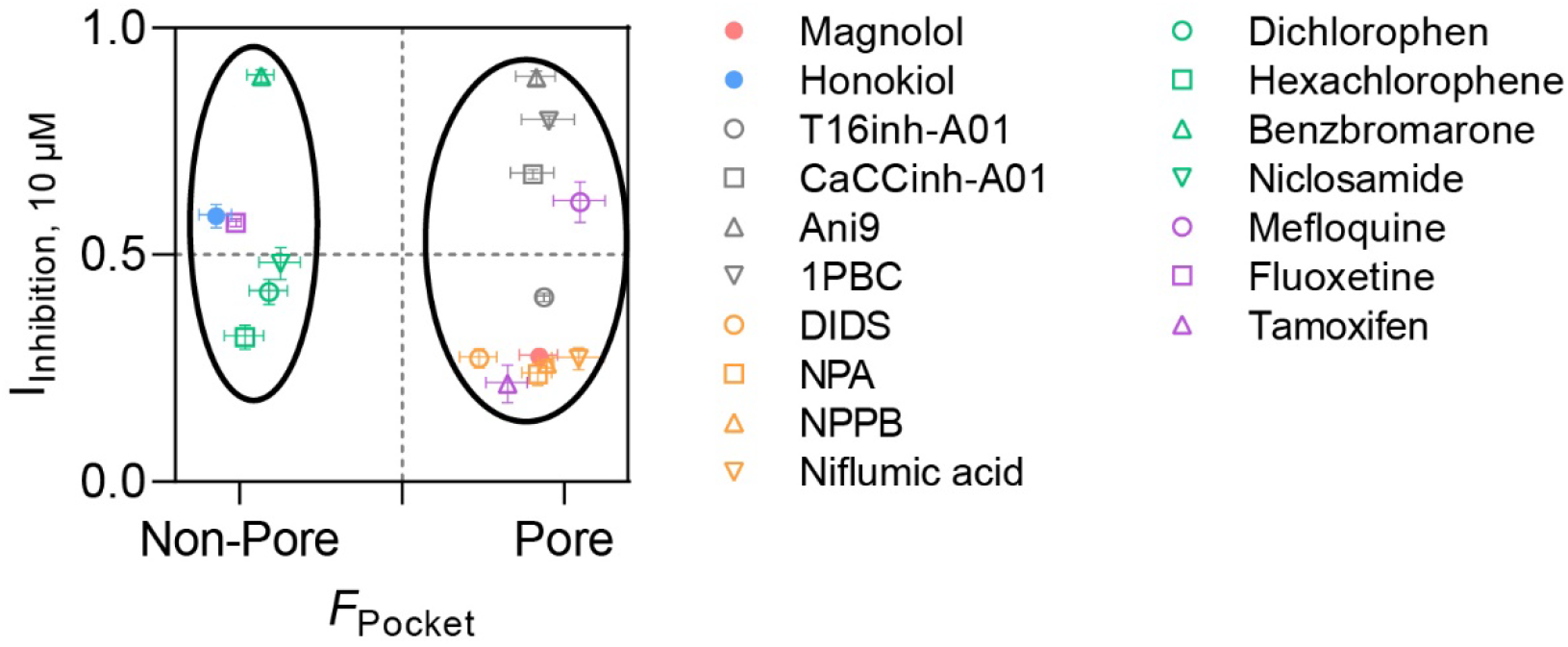
Classification of TMEM16A inhibitors to pore or non-pore blocker. Each inhibitor listed on the right side was tested at 10 μM concentration on the extracellular side in the whole-cell patch clamp configuration. Intracellular Ca^2+^ concentration was fixed at 300 nM. The X-axis denoted as ‘*F*_Pocket_’ was determined by calculating the difference of inhibition rate between the WT, R515A (pore pocket mutant), and F602A/F777A (non-pore pocket mutant) (see Supplementary Fig. 23 and Methods). Drugs that bind to either the pore or the non-pore pocket are grouped together by circles, which was calculated by *k*-means clustering algorithm. Data are expressed as mean ± SEM. Source data are provided as Source Data file.

In conclusion, among the 17 drugs tested, 11 were pore blockers and six were non-pore blockers. Given that DIDS, NPA, NPPB, and niflumic acid are ‘classical’ anion channel blockers, where these drugs block almost all types of anion channels with low potency^1^, they may provide no practical help for new TMEM16A drug designs. With these four drugs excluded, seven drugs were categorized into pore blockers and non-pore blockers included six drugs, which emphasizes the structural diversity among the non-pore blockers and their potential for future drug development (Supplementary Fig. 24).

## DISCUSSION

In this study, we identified the plant-derived regioisomeric compounds, magnolol and honokiol, as inhibitors of the CaCC, TMEM16A, that plays a significant role in cellular processes. We utilized electrophysiology, unbiased molecular docking and clustering, alanine scanning mutagenesis, and MD simulations to identify the binding sites for magnolol and honokiol, and termed them as pore and non-pore pockets, respectively. Interestingly, pore blocker-interacting residues were also crucial for Cl^−^ permeation, whereas non-pore blocker-binding caused structural disruption of the pore pocket residues. Finally, we classified various currently known TMEM16A inhibitors into pore or non-pore blockers and found that a substantial proportion of drugs targeted the non-pore pocket.

Our algorithm to identify drug-binding sites in TMEM16A successfully predicted binding sites for magnolol and honokiol, as well as 1PBC and CaCCinh-A01 (Fig. 2, Supplementary Figs. 1 and 2). Because the three-dimensional surfaces of channel proteins are often very complex, there could be numerous locally stable docking conformations for small ligands whose molecular weight are usually <500 g/mol^28, 38^. We overcame this problem by increasing the number of Vina runs. We then gathered the docking conformations by the channel geometry and clustered them using the *k*-means algorithm (Fig. 2). This process is not limited to Vina or TMEM16A but can be applied to all available docking programs and ion channels. Identifying the drug-binding site of a protein is important because it can aid in rational drug development and lead to development of new and more potent drug with better efficacy.

During our alanine scanning mutagenesis experiments, we found that some mutants of the outer pore region, such as R515A, K603A, E623A, and E633A, showed decreased whole-cell current levels compared to WT TMEM16A (Supplementary Fig. 19). Interestingly, these mutants exhibited a decreased response to the pore blocker (Fig. 3a). This led us to hypothesize that residues that are important for Cl^−^ permeation may also be crucial for interaction with pore blockers. Indeed, when fitted with the ion permeation model developed by the Dutzler group^34^, we successfully analyzed the function of these residues as well as their electrostatic relationships (Figs. 5, 6).

Our findings on electrostatic interactions in the outer pore region identified residues important for TMEM16A function: R515 and K603. These residues have been earlier identified to be important for anion selectivity^13, 22^, which is in line with our results that these residues may directly interact with the permeating anion. Moreover, we observed fine-tuning of R515 and K603 by E623 and E633 (Fig. 5); E623A or E633A mutants increased the energy barrier for ion permeation. In addition, non-pore blocker-binding caused TM3 and TM4 to move away from the center of the pore, resulting in disrupted electrostatic interactions in R515, K603, E623, and E633 (Figs. 7 and 9a). Therefore, the pore and non-pore blockers modulate outer pore residues in a different manner, yet their ultimate result is the same: TMEM16A inhibition (Fig. 9a). It is worthwhile to note that E623 has been shown to be the key residue for extracellular pH sensing in TMEM16A^39^. This further highlights the physiological importance of outer-pore residues. In addition, the outer-pore residues are absolutely conserved among TMEM16A, TMEM16B, and TMEM16F (Supplementary Fig. 25). Therefore, electrostatic interactions in the outer pore region may be an evolutionarily conserved feature in TMEM16 ion channel proteins.

**Fig. 9.**
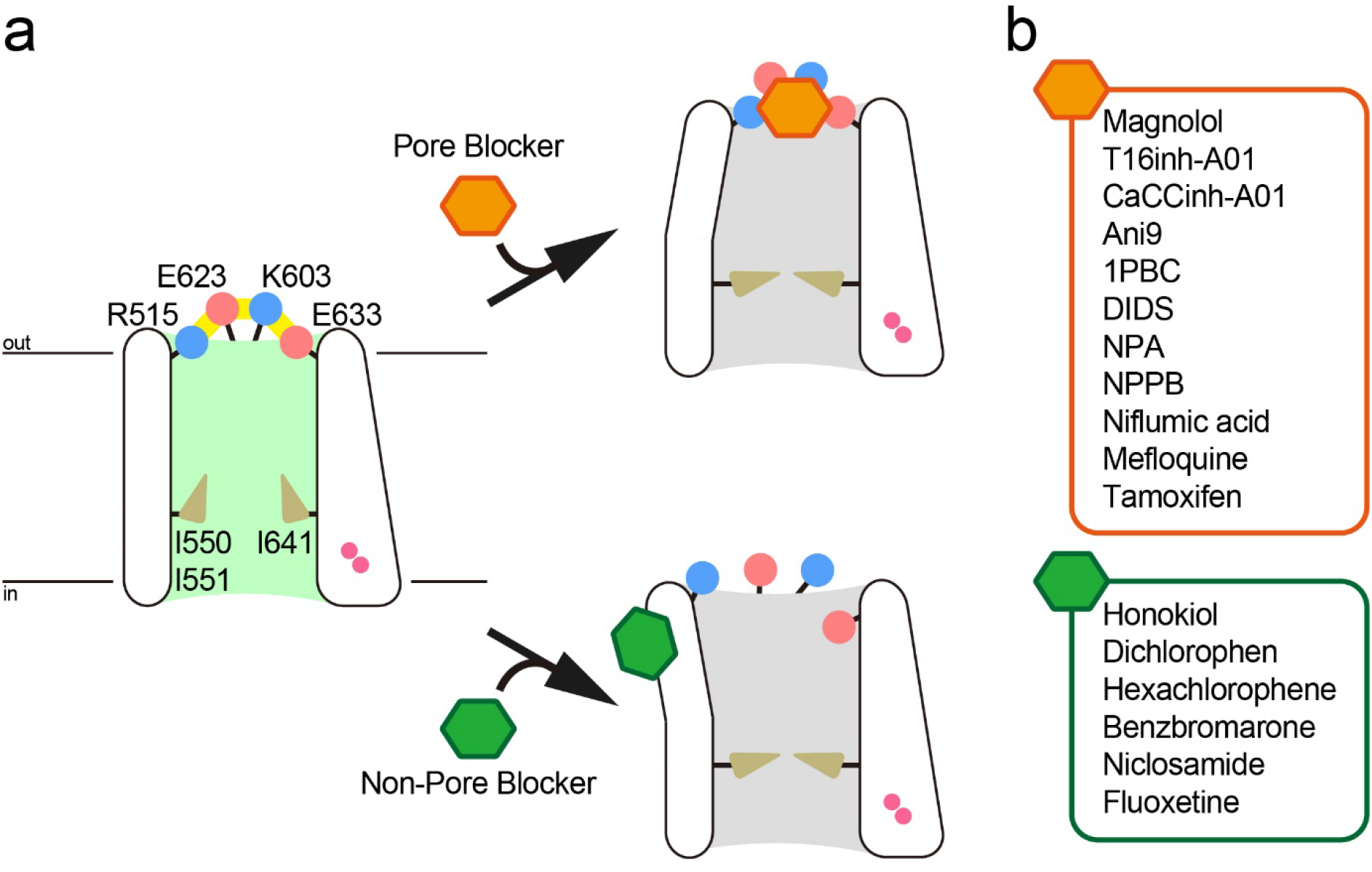
Graphical illustration of TMEM16A blockers. **a.** Pore region in TMEM16A. For the sake of simplicity, only one subunit is shown. The side chains of the positively charged residues R515 and K603 are shown as blue circles, and the side chains of the negatively charged residues E623 and E633 are shown as red circles. The neutral side chains of I550, I551, and I640 are indicated by brown triangles. The two Ca^2+^ ions are indicated by magenta circles. The salt bridges formed by R515, K603, E623, and E633 are indicated by the yellow lines. The conductive pores of TMEM16A are colored in lime, whereas the non-conductive pores are colored in gray. The pore blocker is shown as an orange hexagon, and the non-pore blocker is shown as a green hexagon. The binding regions of pore and non-pore blockers are shown according to their relative locations. **b.** Drugs classified in Fig. 8. The pore blockers are shown in the orange box and the non-pore blockers are shown in the green box.

One study published a binding site for honokiol on TMEM16A, which was different from ours^40^. They identified honokiol binding sites by comparing the sequence differences between TMEM16A and TMEM16B because honokiol was more potent on TMEM16A than TMEM16B. They proposed that R429, K430, and N435 are honokiol-binding sites (Supplementary Fig. 26a). Surprisingly, when we constructed R429A, K430L, and N435G mutants and tested the effects of honokiol, we were unable to observe a difference in the remaining current ratio of the mutants compared to the WT (Supplementary Fig. 26b, c). Moreover, R429, K430, and N435 did not form a 3D pocket, rather protruded out in different directions (Supplementary Fig. 26a). This led us to conclude that these residues are unlikely to harbor an honokiol-binding site in TMEM16A. It is also worth noting that R429 and K430 are located at the PIP_2_-binding site^16^ and near the third Ca^2+^-binding site^19, 30^. In our experiments, the R429A and K430L mutants showed different deactivation kinetics than the WT (Supplementary Figs. 19 and 26d).

Classification of TMEM16A inhibitors based on their binding properties may provide insights into drug-binding pockets in TMEM16 families. We found that T16inh-A01, CaCCinh-A01, and Ani9, which were identified by a large library screening^20, 21^, favorably bound to the pore pocket. Recent identification of the CaCCinh-A01-binding site in the outer pore region is consistent with our data^25^. In addition, DIDS, NPA, NPPB, and niflumic acid, all of which are classical anion channel pore blockers, also favor the pore pocket^36, 37^. Among drugs that bind to the non-pore pocket, niclosamide was recently found to inhibit COVID-19-induced syncytia by blocking TMEM16F^24^. Cryo-EM study with TMEM16F and niclosamide revealed distinct binding site different from our non-pore pocket^41^. Further studies are needed to elucidate the different drug modulation mechanisms of TMEM16 ion channels and scramblases.

In conclusion, our work reveals a novel mechanism for the structural characteristics and pharmacological regulation of TMEM16A and offers a structure-based rational background for future drug development.

## METHODS

### Cell culture and plasmid transfection

HEK293T cells (American Type Culture Collection, USA) were cultured in high-glucose Dulbecco’s modified Eagle’s medium (Thermo Fisher Scientific, USA) supplemented with 10% fetal bovine serum (Thermo Fisher Scientific) and 1% penicillin/streptomycin (Thermo Fisher Scientific). The cells were cultured in a humidified incubator at 37 °C with 20% O_2_ and 5% CO_2_, and were subcultured every 2–3 days in T25 flasks.

The coding region of human TMEM16A (hTMEM16A) ac isoform was subcloned into the pEGFP-N1 mammalian expression plasmid by replacing the EGFP region (pCMV-hTMEM16A) as described previously^42^. Mutations were introduced by PCR-based site-directed mutagenesis method using the Quick Change II XL Site-Directed Mutagenesis Kit (Agilent Technologies, USA) and verified by sequencing (Supplementary Table 2).

Plasmids were transiently transfected into HEK293T cells using Lipofectamine LTX transfection reagent (Thermo Fisher Scientific) according to the manufacturer’s recommendations. HEK293T cells were seeded in 12-well plates and transiently transfected with 1000 ng of total DNA for electrophysiology recordings. Experiments were performed within 24–48 h of transfection.

### Solutions

Whole-cell patch clamp experiment was conducted using a basal extracellular solution containing 140 mM NaCl, 10 mM HEPES, and 2 mM MgCl_2_, with pH adjusted to 7.4 using NaOH, and osmolarity adjusted to ∼310 mOsm with an appropriate amount of sorbitol. The basal pipette solution with 300 nM free Ca^2+^ contained 133bb.4 mM NaCl, 10 mM HEPES, 5 mM EGTA, and 3.3 mM CaCl_2_, 3 mM Mg-ATP, with pH adjusted to 7.2 using NaOH, and osmolarity adjusted to ∼290 mOsm with an appropriate amount of sorbitol. Free Ca^2+^ was calculated using the WEBMAX-C software (C. Patton, Stanford University; https://somapp.ucdmc.ucdavis.edu/pharmacology/bers/maxchelator/webmaxc/webmaxcS.htm).

For the inside-out patch clamp recording to calculate the ion permeation model of TMEM16A^34^, the basal bath solution contained 140 mM NaCl, 10 mM HEPES, 5 mM EGTA for Ca^2+^-free solution; and 128 mM NaCl, 10 mM HEPES, 5 mM EGTA, and 6 mM CaCl_2_ to obtain a 1 mM free Ca^2+^ concentration (both solutions were adjusted to pH 7.2 using NaOH). A 1 mM free Ca^2+^ solution was also used as the pipette solution to maintain uniform ionic conditions.

### Electrophysiology

Transfected cells were transferred to a bath perfused at 5 mL/min and mounted on the stage of an inverted microscope (Nikon, Japan) equipped with a high-density mercury lamp light source for green fluorescence excitation. Microglass pipettes (World Precision Instruments, USA) were fabricated using a PP-830 single-stage glass microelectrode puller (Narishige, Japan), with a resistance of 2–5 MΩ. The liquid junction potential was rectified using an offset circuit prior to each recording. Currents were recorded using an Axopatch 200B amplifier (Molecular Devices, USA) and Digidata 1440A interface (Molecular Devices), digitized at 10 kHz, and low-pass filtered at 5 kHz using pClamp software 10.7 (Molecular Devices). All recordings were performed at room temperature (22–25°C). The whole-cell patch clamp configuration was verified by measuring the series resistance to <10 MΩ, which was compensated before each recording. The inside-out patch-clamp configuration was achieved by briefly exposing the giga-ohm-pipette to air, followed by gentle submerging in the bath solution.

In the whole-cell configuration, TMEM16A currents were recorded at a holding potential of – 60 mV, and drugs were applied after >5 min of break-in to reduce the effects of channel rundown^43^. The drugs were tested for >5 min to obtain a stable current level. To measure the channel activity, cells were held at 0 mV for 0.5 s, then –60, 0, or 60 mV pulse were respectively applied for 1 s, then held at –60 mV for 0.5 s. Channel activity was calculated as the average current density in the last 20 ms of 60 mV pulse.

In the inside-out configuration, cells were initially exposed to free Ca^2+^ solution to examine the background currents, as described previously^34, 44^. The bath solution was then switched to the 1 mM Ca^2+^ solution, and currents were obtained by applying –100 to 120 mV for 300 ms in 20 mV increments. To correct the channel rundown by high Ca^2+^ ^13, 17^, a pulse of 80 mV was applied for a 50 ms before each voltage jump. The obtained currents were initially normalized at an 80 mV pre-pulse and then expressed as normalized current at 120 mV (I/I_+120 mV_).

All chemicals used in electrophysiology were purchased from Sigma-Adrich (St. Louis, MO, USA) except for 1PBC, which was purchased from ChemBridge (San Diego, CA, USA). All experiments were performed in independent biological replicates, and the mounting chamber was replaced with new cells after each drug.

### Ion permeation model of TMEM16A

We used the TMEM16A ion permeation model initially described by Läuger^45^ and modified by Dutzler’s group^34^. This model assumes that multiple energy barriers lie on the ion permeation pathway, and calculation of the energy difference between barriers can determine the pore property of TMEM16A. Although the energy barriers are largely hypothetical and do not necessitate the actual structural presence of such calculations, the success of this model in explaining TMEM16A behavior^33, 46^ encouraged us to examine and utilize it in our experiments.

The normalized IV relationship acquired from the inside-out patch clamp configuration was fitted using the following equation:

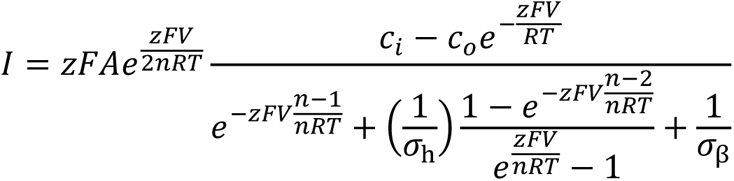

In this equation, *z* is the charge of the permeating ion, which in our experiments was set to – 1; *F* is the Faraday constant; *R* is the gas constant; *T* is the absolute temperature; *c_i_* and *c_o_* are intracellular and extracellular concentrations of permeating ions; *n* is the number of energy barriers, which was previously calculated as 3^34^; *σ*_β_ is the rate constant for outward movement of permeating ions from the innermost to the outermost barrier; and *σ*_h_ is the rate constant from the middle to the outermost barrier. Because *σ*_β_ and *σ*_h_ are rate constants, they can be used to calculate the energy difference between the barriers as follows:

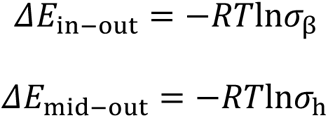

In our experiments, we mainly conducted mutagenesis experiments in the outer pore region of TMEM16A; thus, the above equation can be modified to set the reference energy level of the innermost barrier.

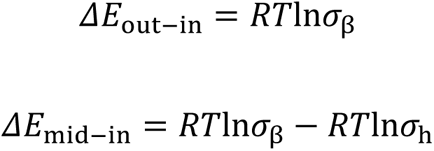

### Double mutant cycle analysis

Double mutant cycle analysis can be used to assess the energy coupling of two residues in the protein^35, 47, 48^. In the present case, the energy state can be calculated from wild-type protein (WT), two single mutants (Mut A and Mut B), and the corresponding double mutant protein (Mut AB) channels, as described in the equations below (Supplementary Fig. 27). The coupling energy can be calculated as follows: The total difference in the free energy perturbation by the mutants must be the same, regardless of the pathway taken^48^.

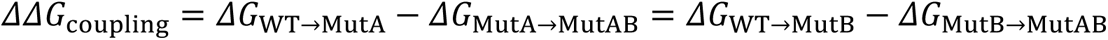

From this equation, we obtain the final equation.

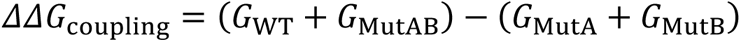

If Mut A and Mut B function independently of one another, the coupling energy should be close to zero. If the two mutants directly interact with each other, the free energy perturbation should deviate from zero. The coupling energy is commonly expressed in absolute values. Two coupling energy values were obtained from the TMEM16A ion permeation model, *E*_mid-in_ and *E*_out-in_.

### Electrophysiology data analysis

The concentration-dependent inhibition rates of drugs were fitted to the Hill equation,

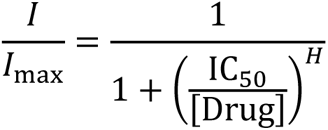

where *I* is the current level at the designated drug concentration, *I*_max_ is the current level before drug treatment, IC_50_ is the half-maximal inhibitory concentration of the drug, and *H* is the Hill coefficient.

To calculate the proportion of drug binding to the pore, and non-pore pockets, we assumed that: (i) The pore pocket mutant (R515A) and the non-pore pocket mutant (F602A/F777A) completely abolished drug binding at 10 μM concentration, and (ii) the pore and non-pore pockets are the only drug-binding sites in TMEM16A. Based on these hypotheses, we calculated the proportion of drug binding in the pore pocket by comparing the difference in inhibition rates between the WT and the non-pore pocket mutant, and vice versa. The calculations are as follows:

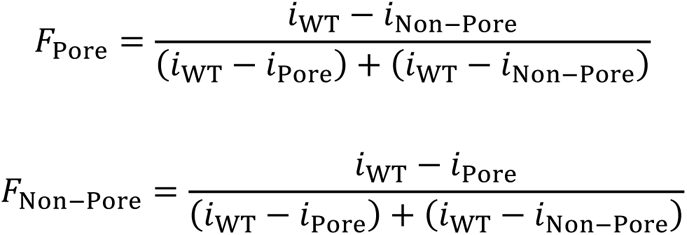

*F*_Pore_ and *F*_Non-Pore_ are the fractions of the drug in each pocket; *i*_WT_, *i*_Pore_, and *i*_Non-Pore_ are the remaining proportion of current for the drug in the WT, pore pocket mutant (R515A), or non-pore pocket mutant (F602A/F777A) respectively, at 10 μM drug concentration. Note that *F*_Pore_ + *F*_Non-Pore_ = 1, which corresponds to our second hypothesis.

### Unbiased molecular docking and clustering

AutoDock Vina^28^ was used for molecular docking throughout the study. The ‘global’ docking, where the entire TMEM16A was located within the Vina grid, was initially performed. The ‘exhaustiveness’ parameter was set to 24, ensuring sufficient calculation for identifying the drug binding poses^28^. The overall grid size was 150 × 150 × 150 Å^3^.

Initially, molecular docking was performed by placing the entire TMEM16A structure in the Vina grid with 150 × 150 × 150 Å^3^ space. However, even increasing the ‘exhaustiveness’ parameter to 200, where the default value is 8, reproducible docking poses between each run could not be generated. We assumed that this was due to the complex surface structure of TMEM16A, since each run in Vina was thought to find only one of many ‘locally stable’ docking poses in the whole structure^28^. To overcome this problem, we increased the number of Vina runs. Thus, 100 independent Vina runs were performed, and 10 docking poses were generated in each run, creating 1,000 docking poses.

To remove false-positive docking results and identify significant drug-binding sites, docking poses were clustered based on their population. Initially, the center of mass for each docking pose was calculated. Because TMEM16A or TMEM16F exists as a homodimer, center of masses in the ‘B’ subunit (referred to as the left subunit in Fig. 2a and Supplementary Figs. 1 and 2) were moved to the corresponding positions in the ‘A’ subunit (referred to as the right subunit in Fig). Thus, a rotation matrix along the Z axis was used for inter-subunit relocation.

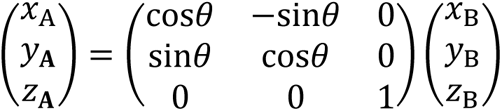

*x*_A_ or *x*_B_, *y*_A_ or *y*_B_, and *z*_A_ or *z*_B_ refer to 3-dimensional coordinates of each subunit, and *θ* is the rotation angle, which was 180° for the TMEM16A dimer.

The *k*-means clustering algorithm was used to cluster the docking poses based on the 3-dimentional coordinates of the A subunit. Assuming that the number of clusters was set to *n*, we defined ‘major’ clusters as clusters with greater than or equal to average number of binding poses (≥1000/*n*), and ‘minor’ clusters as clusters with less than average number of binding poses (<1000/*n*). The number of clusters was initially set to 3 and increased by 1 until all major clusters contained more than 95% of docking poses within a 25 × 25 × 25 Å^3^ space. Supplementary Table 1 summarizes the overall clustering results for magnolol and honokiol from Fig. 1.

After identification of the pore and non-pore pockets, pocket-specific docking was performed in a 25 × 25 × 25 Å^3^ space. The exhaustiveness value was set to 24, and several independent dockings were performed to ensure that the docking poses were identical between each run.

### Molecular dynamics simulations

The cryo-EM structure of the Ca^2+^-bound mouse TMEM16A (PDB ID 5OYB) was used as the template for human TMEM16A in the SWISS-MODEL server^49, 50^. The overall sequence identity was 92.28%, GMQE score was 0.67, and QMEAN score was –4.24. Several missing short loops in the cryo-EM structure (T260-M266, L467-F487, and L669-K682) were modelled using the SWISS-MODEL server. The missing long loops, including the N- and C-termini (M1-P116, Y131-V164, E911-L960), were presumably dynamic and thus not included in model building. To generate the Ca^2+^-bound human TMEM16F structure, Ca^2+^-bound mouse TMEM16F (PDB ID: 6QP6) was used. The same procedures were conducted with the SWISS-MODEL server as for TMEM16A.

The human TMEM16A structure was initially embedded in the 1-palmitoyl-2-oleoylphosphatidylcholine (POPC) lipid bilayer using a CHARMM-GUI web server^51, 52^. For simulation of the drug-bound TMEM16A, magnolol and honokiol were docked in the pore and non-pore pockets respectively, in both subunits of TMEM16A before POPC bilayer insertion. The system was then neutralized and solvated in a 150 mM NaCl environment. The final system contained approximately 570 phospholipid molecules in a ∼150 × 150 × 170 Å^3^ box, with a total of ∼360,000 atoms. All simulations were performed in the Amber force field^53^ and CUDA-enabled Amber20^54^. The parameters for magnolol and honokiol were assigned using the AnteChamber module in Amber20. The particle mesh Ewald algorithm was used to evaluate the long-distance electrostatic interactions. The van der Waals interactions for a smoothing function were considered to be between 10 and 12 Å. The system was minimized for 5000 steps using the steepest descent algorithm. Six equilibrium steps were applied to the protein and membrane, and the restraints were gradually reduced to zero. Complete electronic evaluation was performed every 2 fs using the SHAKE algorithm^55^. The pressure and temperature were respectively maintained at 1 atm and 298 K throughout the simulation using the Nosé–Hoover Langevin piston method and Langevin dynamics, respectively^56, 57^. No anisotropic cell fluctuations were observed.

Three independent simulations for drug-free, magnolol-bound, and honokiol-bound TMEM16A were conducted. Each production was run for 600 ns, resulting in a 1.8 μs simulation time for each condition with a total of 5.4 μs. The last 300 ns in each production run were analyzed at 100 ps interval for further examination using VMD^58^.

### Molecular model presentations

HOLE^32^ software was used to calculate the ion-permeating pore region of TMEM16A. Graphical pore data were obtained from HOLE and reconstructed in ChimeraX^59^ for better visualization. All graphical illustrations of TMEM16A structures were created using ChimeraX^59^. The 2D structure of the TMEM16A inhibitors was drawn using the RDKit (https://www.rdkit.org/) module in Python.

### Statistical analysis

The data analysis was performed using Clampfit 10.7 (Molecular Devices), GraphPad Prism 9.4.1 (GraphPad Software, USA), and Excel (Microsoft, USA). For analyzing and fitting the ion permeation model, an in-house Python script using NumPy (https://numpy.org) and SciPy (https://scipy.org) were used. Curve fitting was performed using ‘curve_fit’ function in SciPy, and the Jacobian matrix was used for 95% confidence interval calculations. All data are expressed as mean ± SEM values, except for the estimated parameters, which are expressed as estimated values ± 95% confidence intervals. The standard errors are propagated as follows:

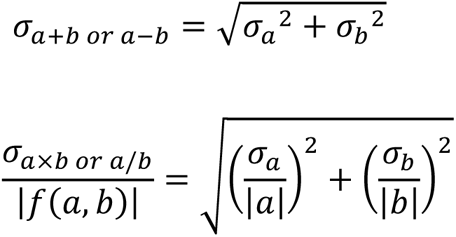

To calculate the energy barriers from the rate constants, the mean (*μ*) and standard deviation (*σ*) of *Y = ln (X)* were calculated using the following equations:

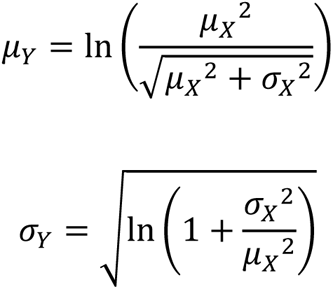

Two-sided unpaired *t*-tests were performed to compare the independent groups. For multiple-group comparisons, one-way analysis of variance (ANOVA) with Dunnett’s post hoc test was used. Statistical significance was set at *p* < 0.05.

## Data Availability

The data that support the findings of this study are available from the corresponding author upon reasonable request. Source data are provided with this article.

## Code Availability

Custom python code for unbiased molecular docking and clustering method for TMEM16A is available at Github (doctroh/TMEM16A-unbiased-molecular-docking-and-clustering).

## Contributions

J.H.N., W.L., and W.K.K. conceived and supervised the study. J.H.N., W.L., W.K.K., and J.W.R. designed experiments. J.W.R. and H.Y.G. provided new tools and reagents. J.W.R. performed molecular docking and clustering, electrophysiology, MD simulation, and data analysis. J.W.R., H.Y.G., W.K.K., W.L., and J.H.N. discussed the results and commented on the manuscript accordingly. J.W.R. wrote and revised the manuscript in consultation with J.H.N., W.L., B.W. and W.K.K.

## ACKNOWLEDGEMENTS

We thank Dr. Byung-Chang Suh (Daegu Gyeongbuk Institute of Science and Technology) and Dr. Sung Joon Kim (Seoul National University) for critical comments on the manuscript. This research was supported by the Bio & Medical Technology Development Program of the National Research Foundation (NRF) funded by the Ministry of Science, ICT & Future Planning (MSIT) (NRF-2018M3A9F3082269 for W.K.K.) and by the Basic Science Research Program through an NRF grant funded by MSIT (No. 2021R1A2C1004344 for W. L.). This research was supported by a grant of the MD-Phd/Medical Scientist Training Program through the Korea Health Industry Development Institute (KHIDI), funded by the Ministry of Health & Welfare, Republic of Korea (for J.W.R.).

