## supplementary information for "Distinct modulation of calcium-activated chloride channel TMEM16A by a novel drug-binding site"

Supplementary Figs. 1–27

Supplementary Tables 1–2

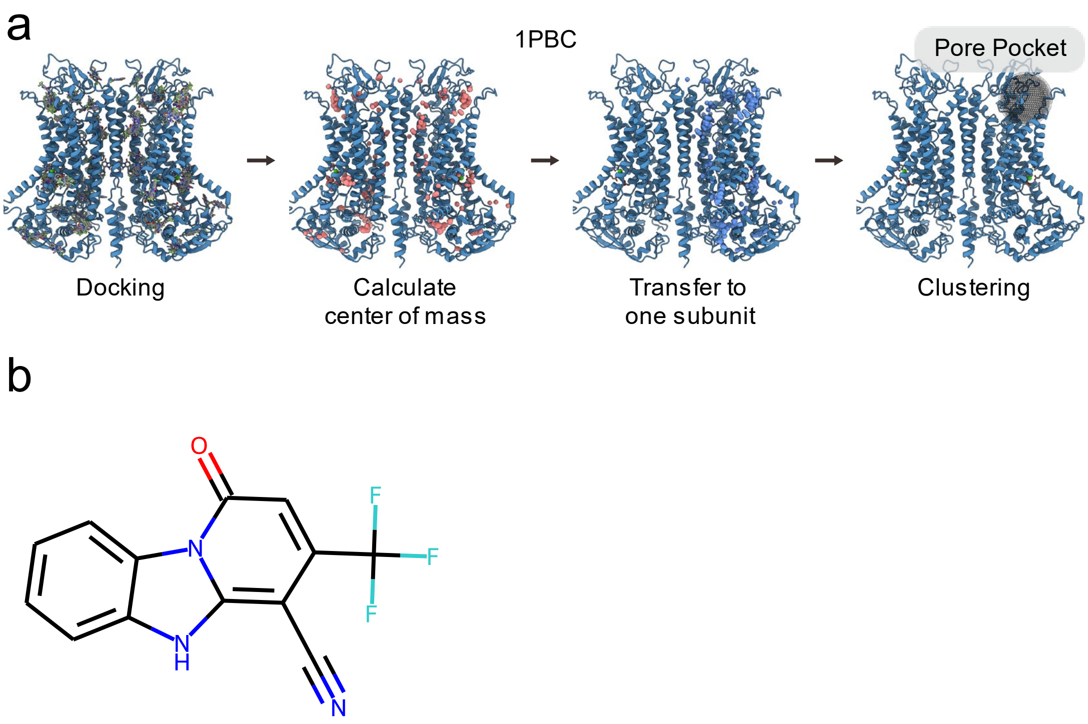


**Supplementary Fig. 1. Unbiased molecular docking and clustering for 1PBC.**

**a.** Unbiased molecular docking and clustering results for 1PBC. **b.** 2D structure of 1PBC. 1PBC: 1-hydroxy-3-(trifluoromethyl)pyrido[1,2-a]benzimidazole-4-carbonitrile.

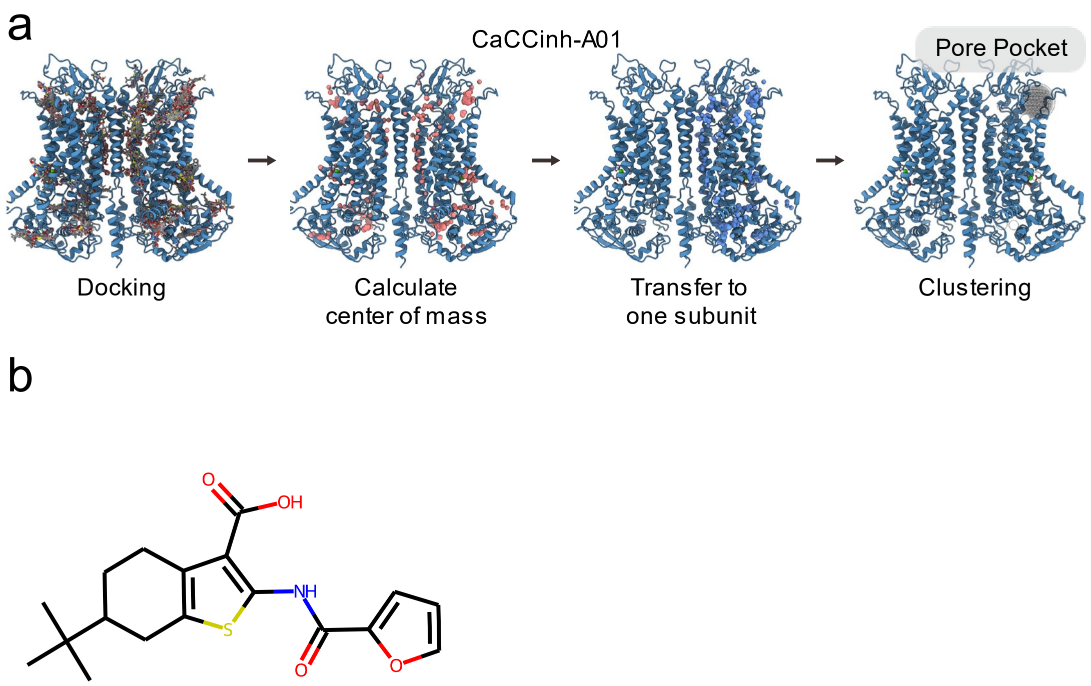


**Supplementary Fig. 2. Unbiased molecular docking and clustering for CaCCinh-A01.**

**a.** Unbiased molecular docking and clustering results for CaCCinh-A01. **b.** 2D structure of CaCCinh-A01.

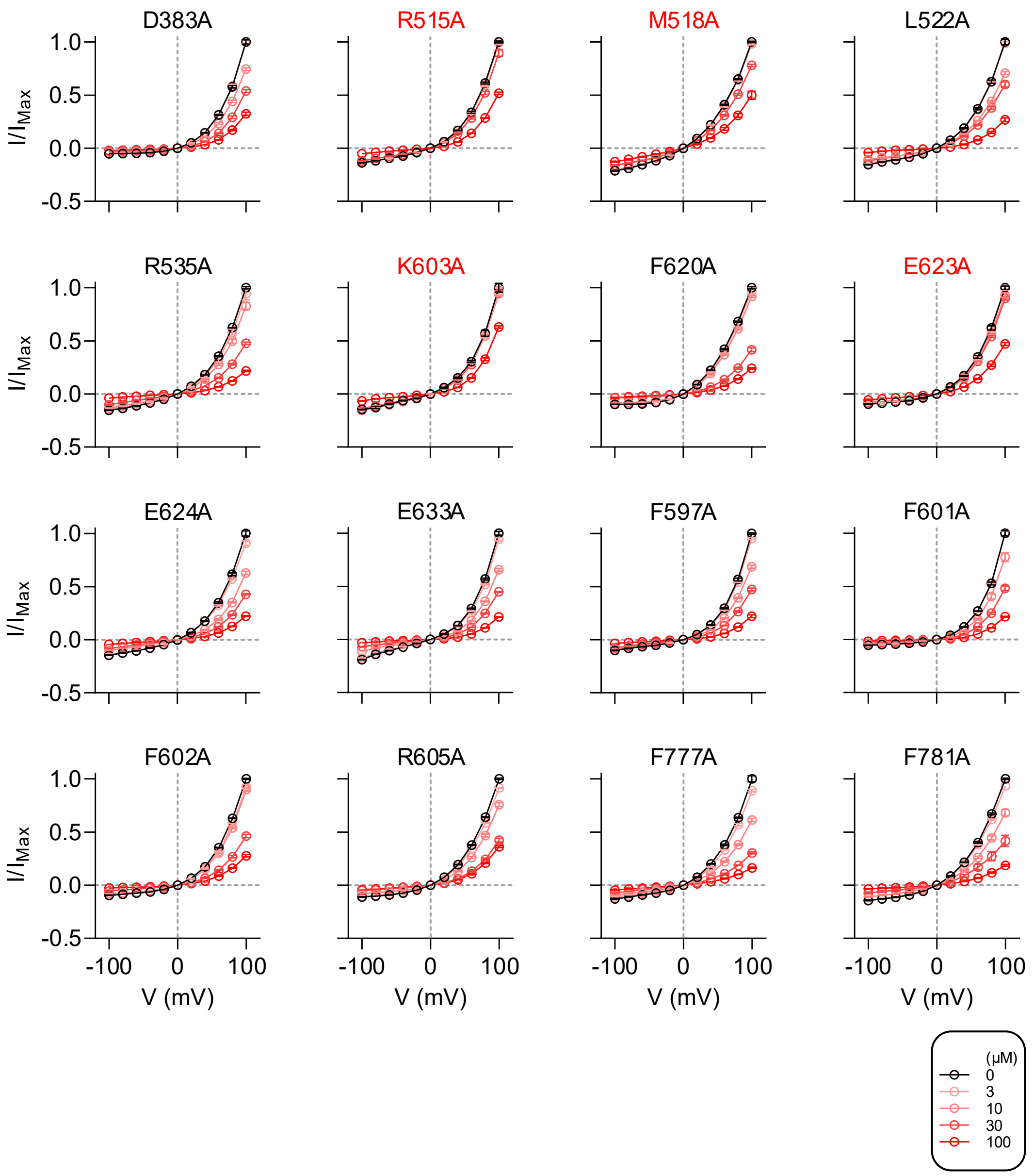


**Supplementary Fig. 3. Current-voltages relationship in alanine-scanning mutagenesis with magnolol administration.**

Current-voltage relationships of TMEM16A whole-cell currents upon magnolol administration on the extracellular side. Intracellular $Ca^{2+}$ was fixed at 300 nM. Concentrations of magnolol are noted in the box. Data are expressed as the mean ± SEM. Source data are provided as Source Data file.

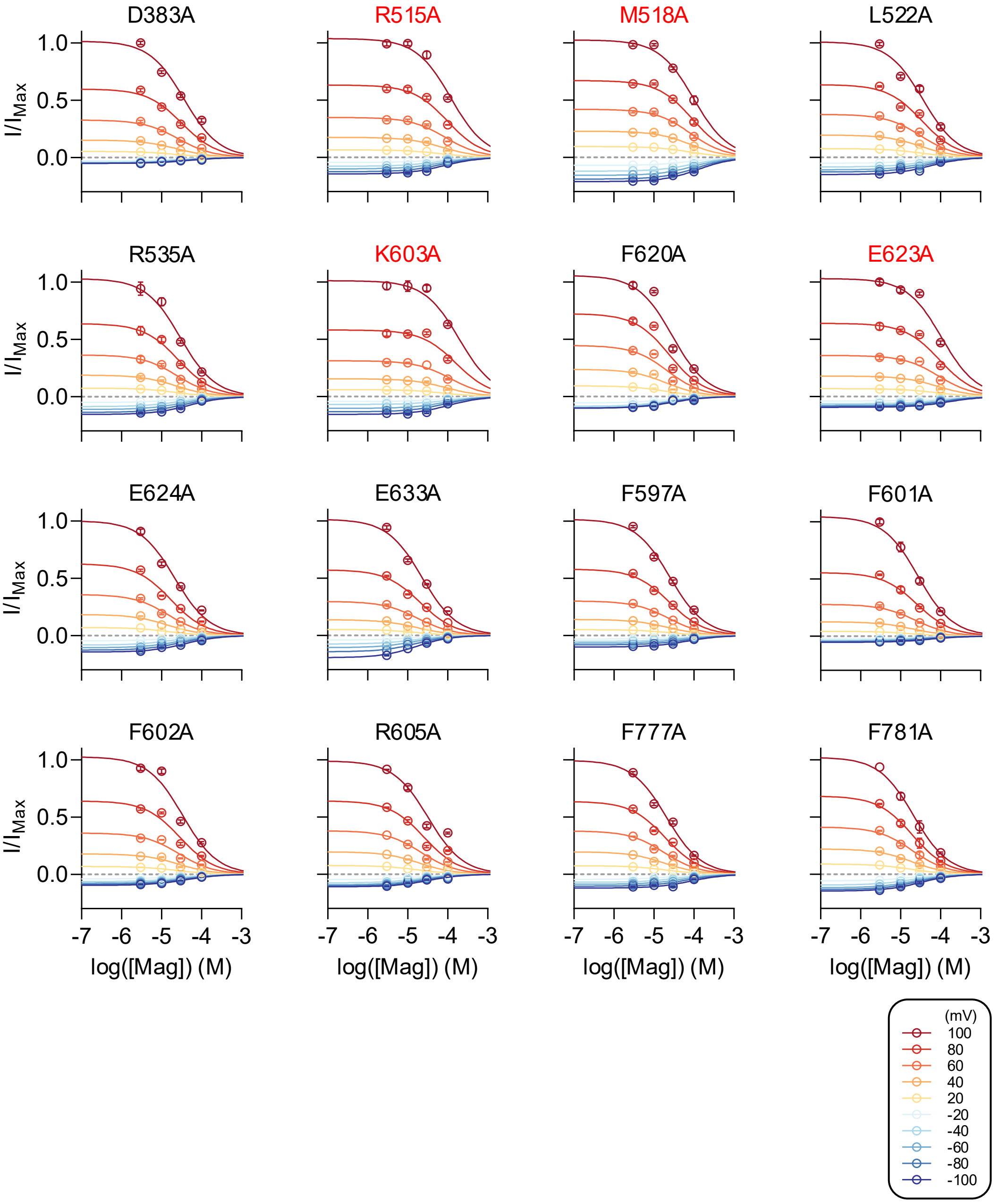


**Supplementary Fig. 4. Dose-response relationships in alanine-scanning mutagenesis with magnolol administration.**

Dose-response relationships of TMEM16A whole-cell currents upon magnolol administration on the extracellular side. Intracellular $Ca^{2+}$ was fixed at 300 nM. Curves were fitted to the Hill equation. Applied voltages are noted in the box. Data are expressed as the mean ± SEM. Source data are provided as Source Data file.

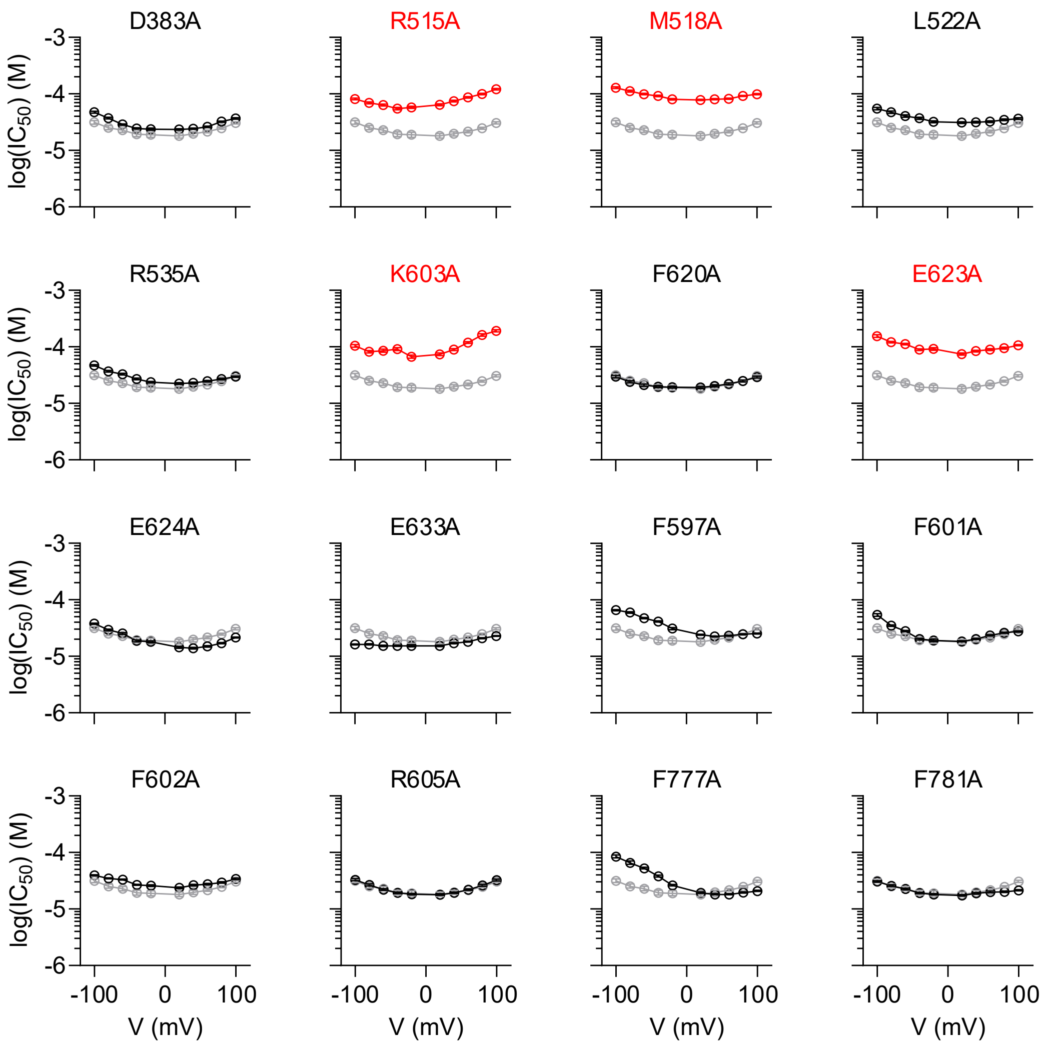


**Supplementary Fig. 5. $IC_{50}$ values in alanine-scanning mutagenesis with magnolol administration.**

$IC_{50}$ values of TMEM16A whole-cell currents upon magnolol administration on the extracellular side. Intracellular $Ca^{2+}$ was fixed at 300 nM. $IC_{50}$ values were calculated by fitting to the Hill equation. Data are expressed as the estimated value ± 95% confidence interval. Source data are provided as Source Data file.

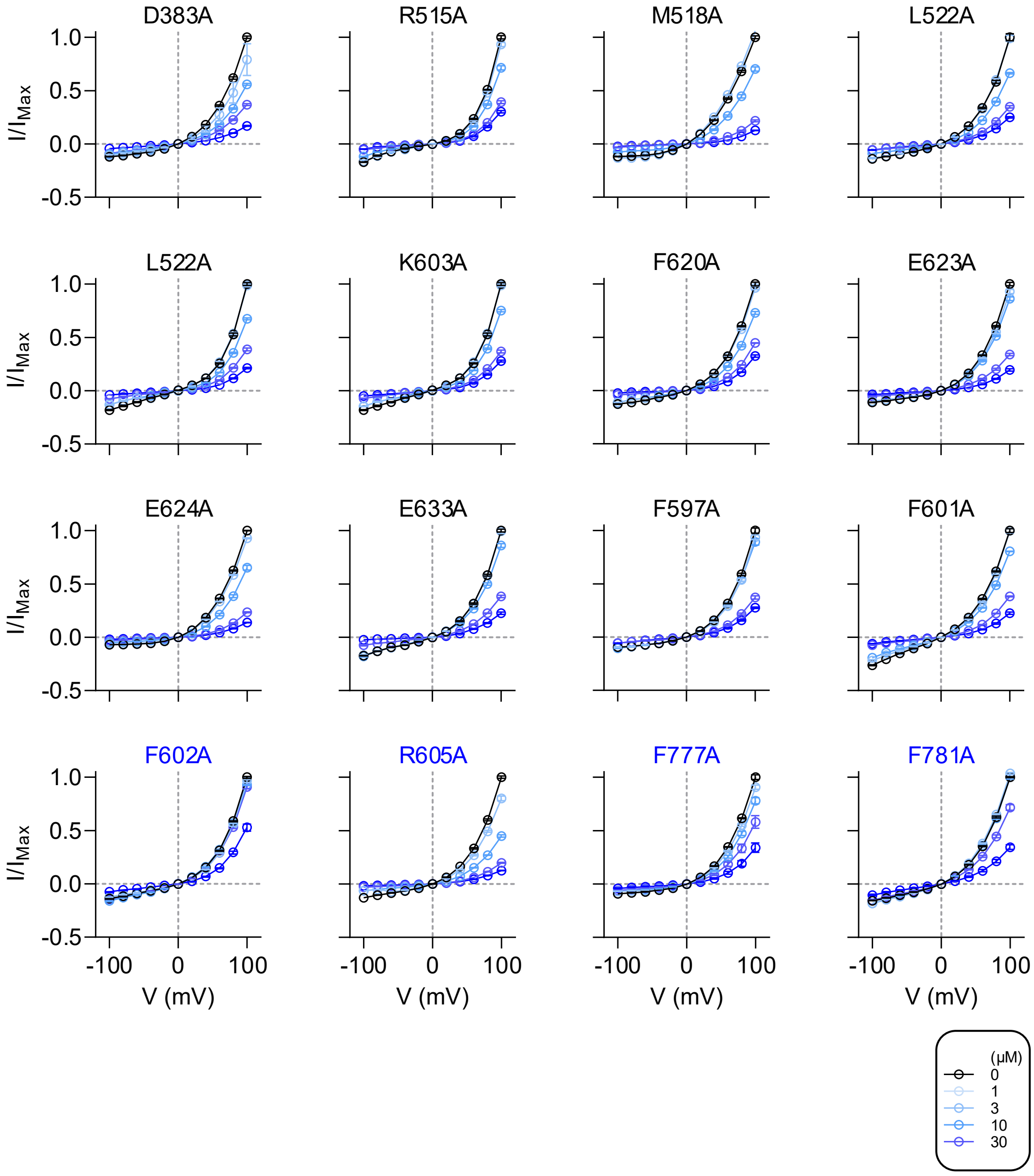


**Supplementary Fig. 6. Current-voltages relationship in alanine-scanning mutagenesis with honokiol administration.**

Current-voltage relationships of TMEM16A whole-cell currents upon honokiol administration on the extracellular side. Intracellular $Ca^{2+}$ was fixed at 300 nM. Concentrations of honokiol are noted in the box. Data are expressed as the mean ± SEM. Source data are provided as Source Data file.

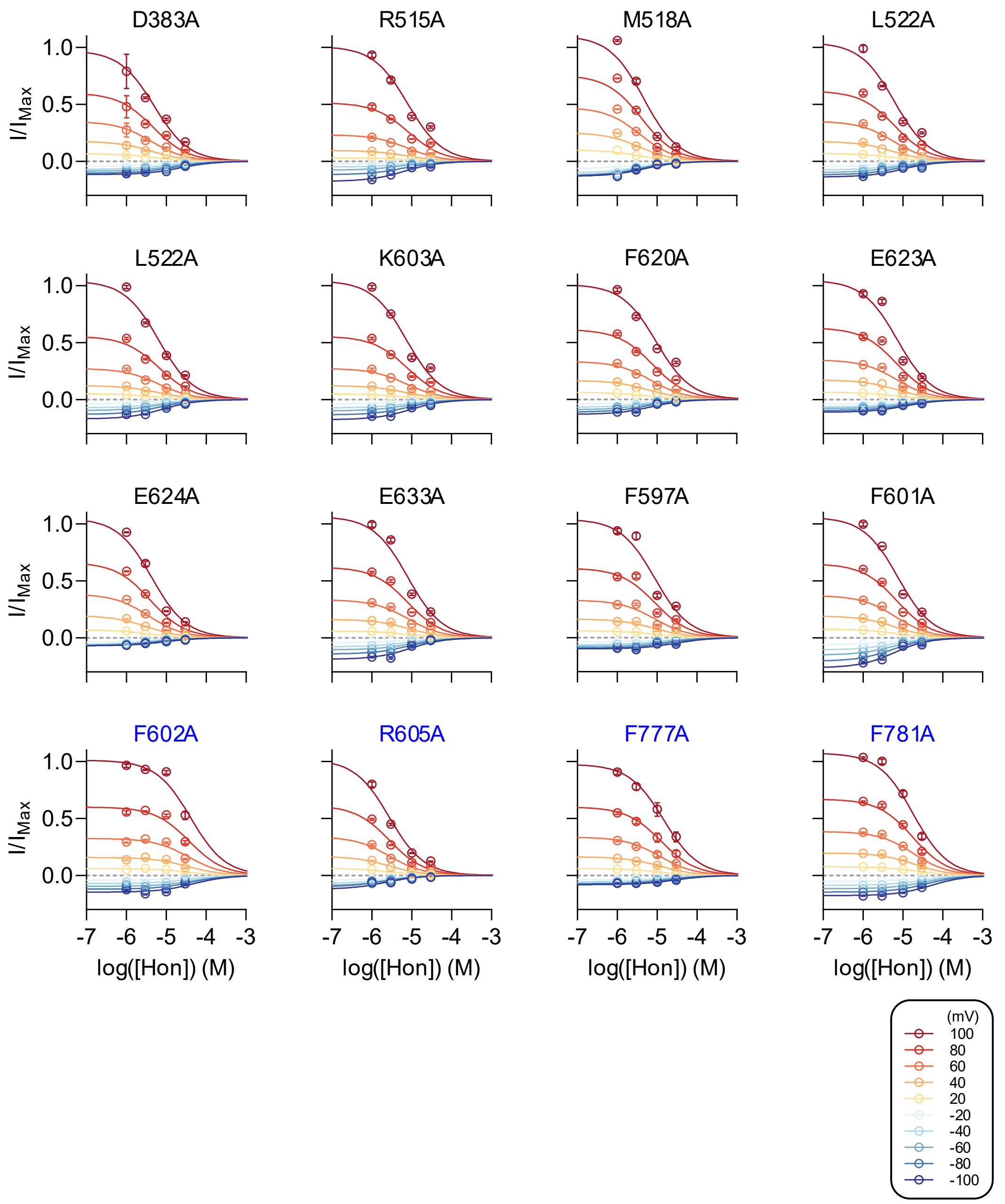


**Supplementary Fig. 7. Dose-response relationships in alanine-scanning mutagenesis with honokiol administration.**

Dose-response relationships of TMEM16A whole-cell currents upon honokiol administration on the extracellular side. Intracellular $Ca^{2+}$ was fixed at 300 nM. Curves were fitted to the Hill equation. Applied voltages are noted in the box. Data are expressed as the mean ± SEM. Source data are provided as Source Data file.

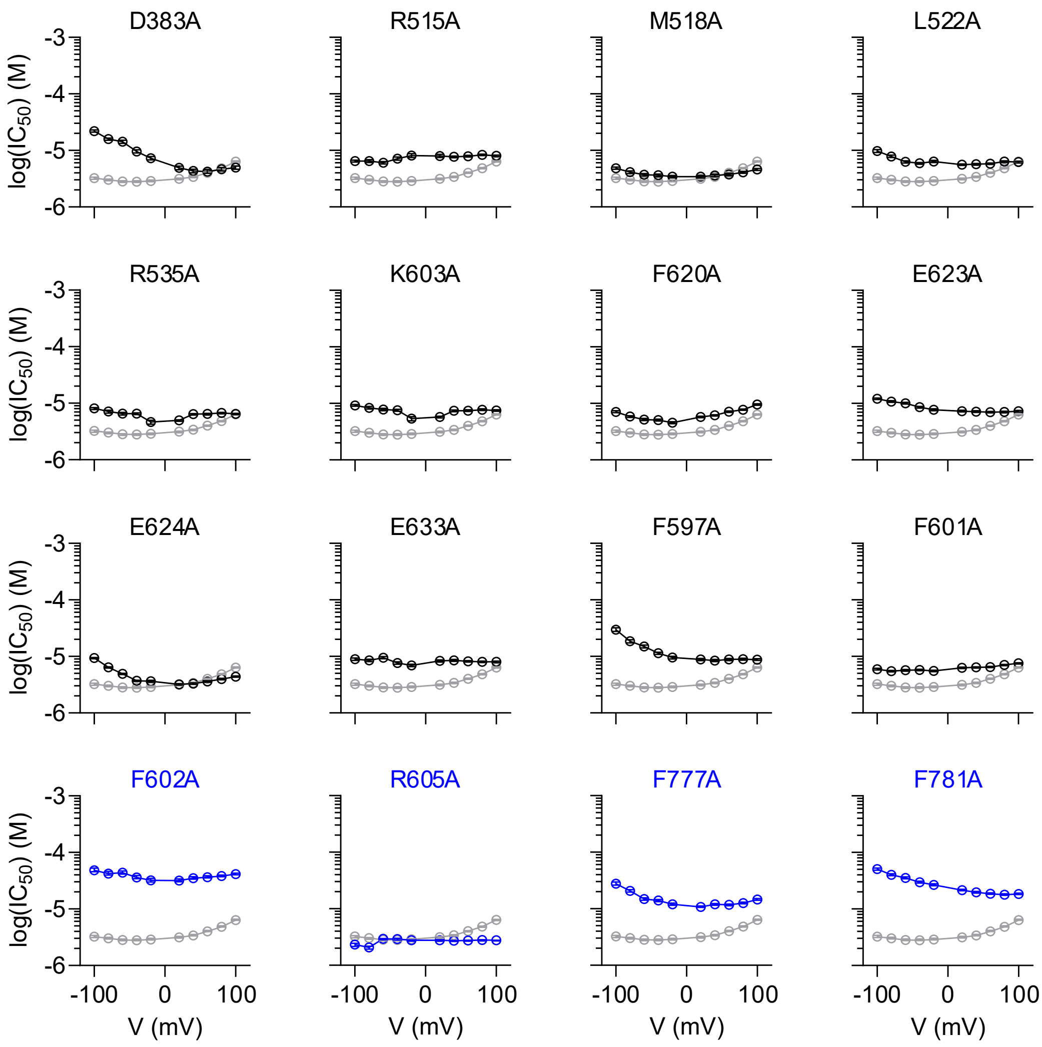


**Supplementary Fig. 8. $IC_{50}$ values in alanine-scanning mutagenesis with honokiol administration.**

$IC_{50}$ values of TMEM16A whole-cell currents upon honokiol administration on the extracellular side. Intracellular $Ca^{2+}$ was fixed at 300 nM. $IC_{50}$ values were calculated by fitting to the Hill equation. Data are expressed as the estimated value ± 95% confidence interval. Source data are provided as Source Data file.

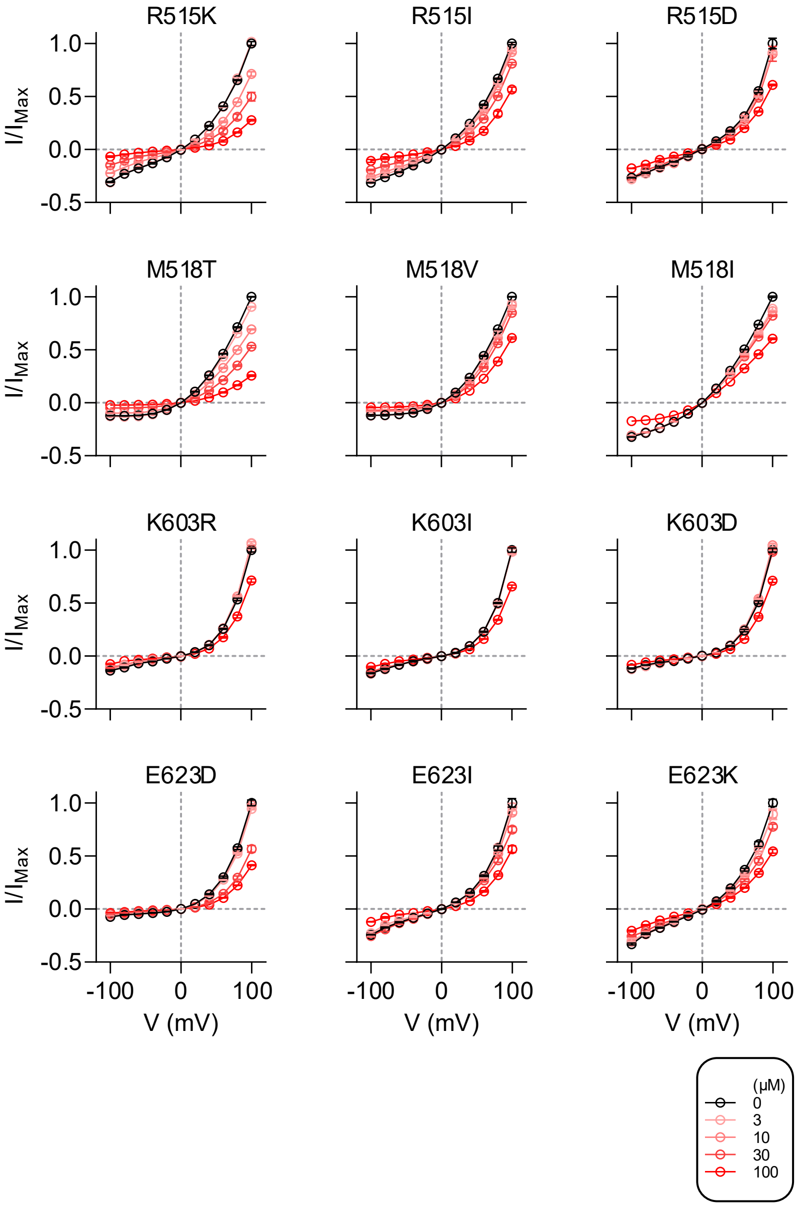


**Supplementary Fig. 9. Current-voltages relationship of side chain mutagenesis on magnolol administration.**

Current-voltage relationships of TMEM16A whole-cell currents upon magnolol administration on the extracellular side. Intracellular $Ca^{2+}$ was fixed at 300 nM. Concentrations of magnolol are noted in the box. Data are expressed as the mean ± SEM. Source data are provided as Source Data file.

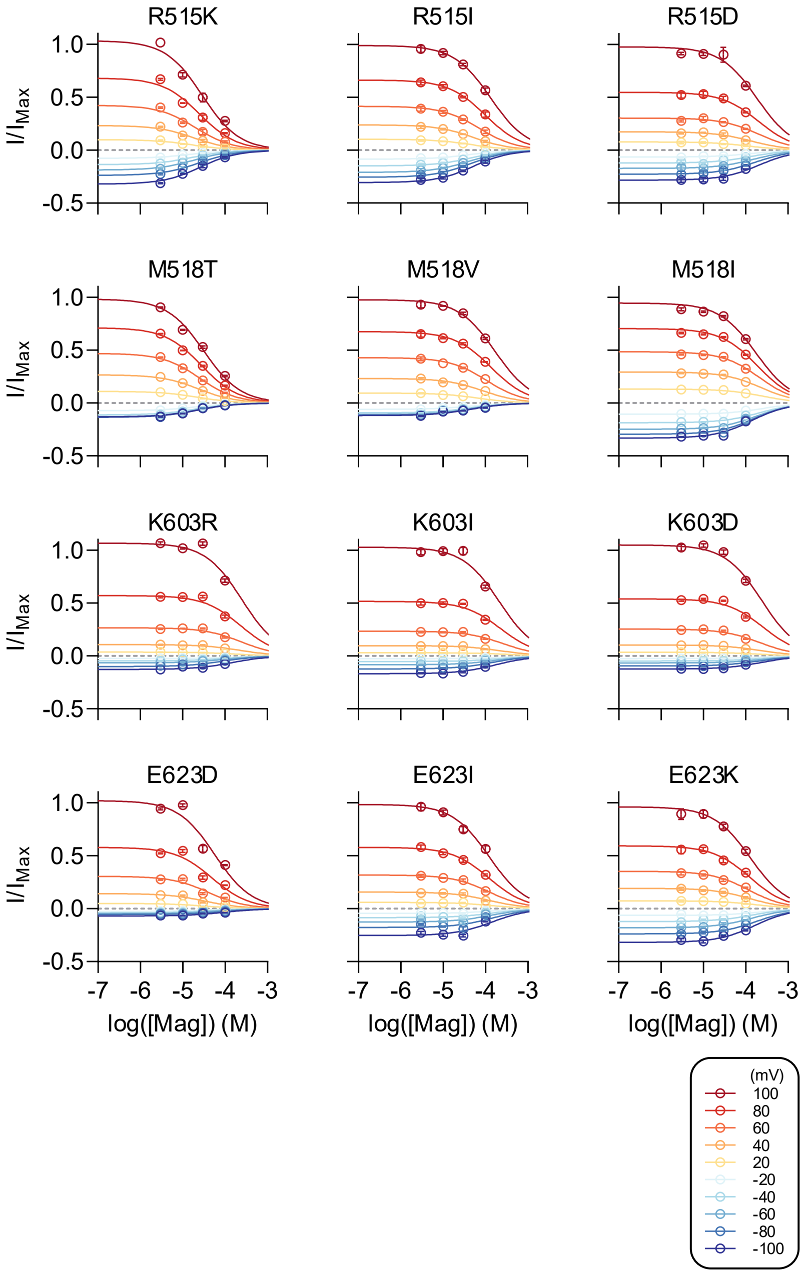


**Supplementary Fig. 10. Dose-response relationships of side chain mutagenesis on magnolol administration.**

Dose-response relationships of TMEM16A whole-cell currents upon magnolol administration on the extracellular side. Intracellular $Ca^{2+}$ was fixed at 300 nM. Curves are fit to the Hill equation. Applied voltages are noted in the box. Data are expressed as the mean ± SEM. Source data are provided as Source Data file.

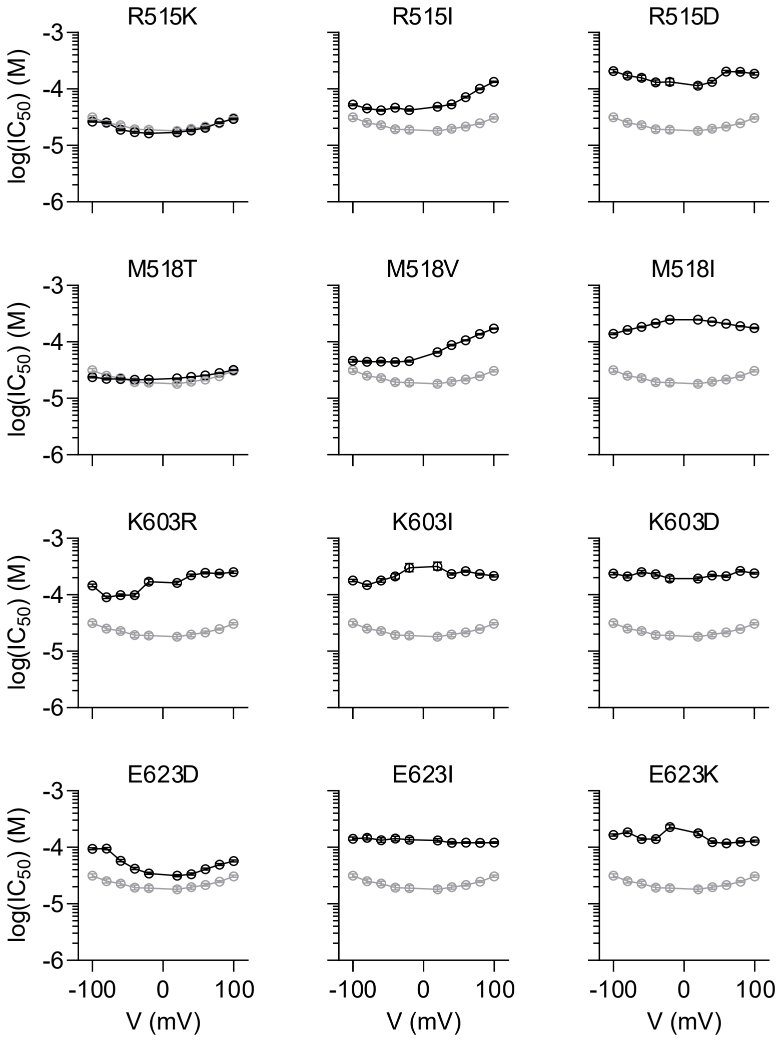


**Supplementary Fig. 11. $IC_{50}$ values of side chain mutagenesis on magnolol administration.** $IC_{50}$ values of TMEM16A whole-cell currents upon magnolol administration on the extracellular side. Intracellular $Ca^{2+}$ was fixed at 300 nM. $IC_{50}$ values were calculated by fitting to the Hill equation. Data are expressed as the estimated value ± 95% confidence interval. Source data are provided as Source Data file.

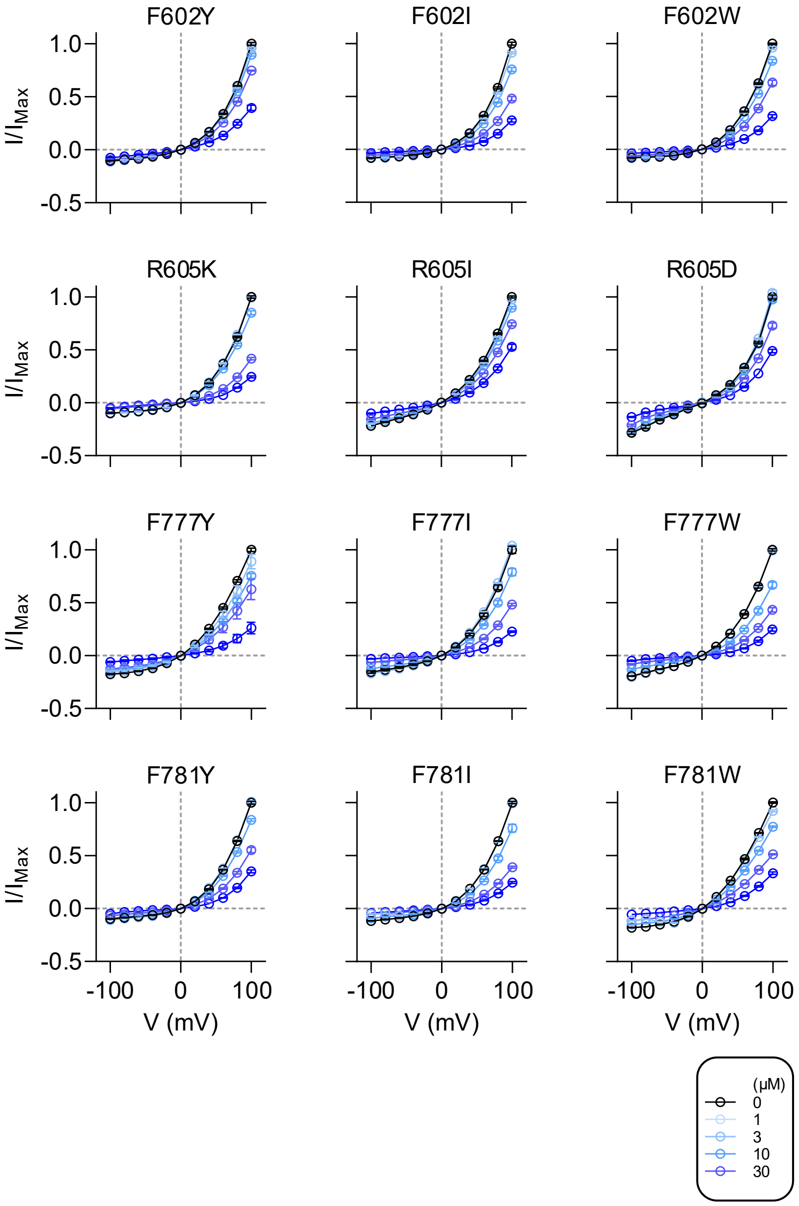


**Supplementary Fig. 12. Current-voltages relationship of side chain mutagenesis on honokiol administration.**

Current-voltage relationships of TMEM16A whole-cell currents upon honokiol administration on the extracellular side. Intracellular $Ca^{2+}$ was fixed at 300 nM. Concentrations of honokiol are noted in the box. Data are expressed as the mean ± SEM. Source data are provided as Source Data file.

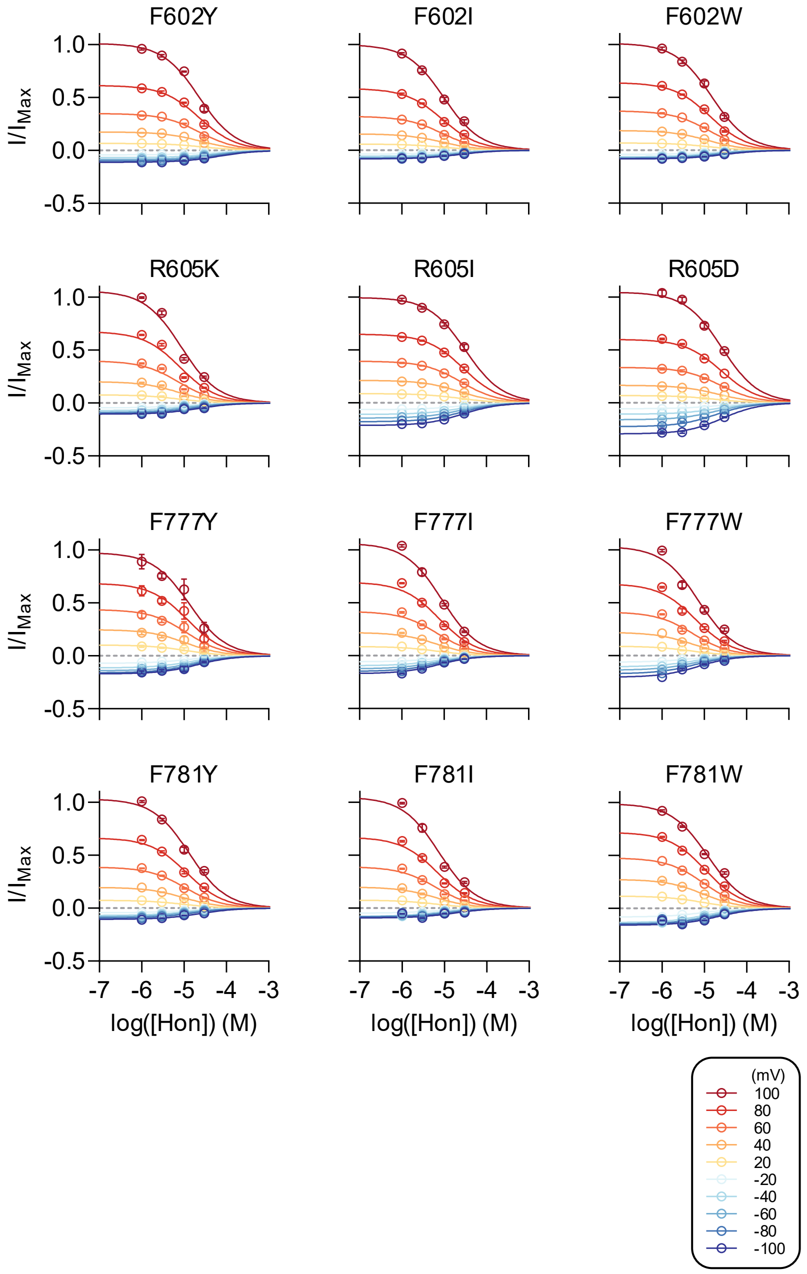


**Supplementary Fig. 13. Dose-response relationships of side chain mutagenesis on honokiol administration.**

Dose-response relationships of TMEM16A whole-cell currents upon honokiol administration on the extracellular side. Intracellular $Ca^{2+}$ was fixed at 300 nM. Curves are fit to the Hill equation. Applied voltages are noted in the box. Data are expressed as the mean ± SEM. Source data are provided as Source Data file.

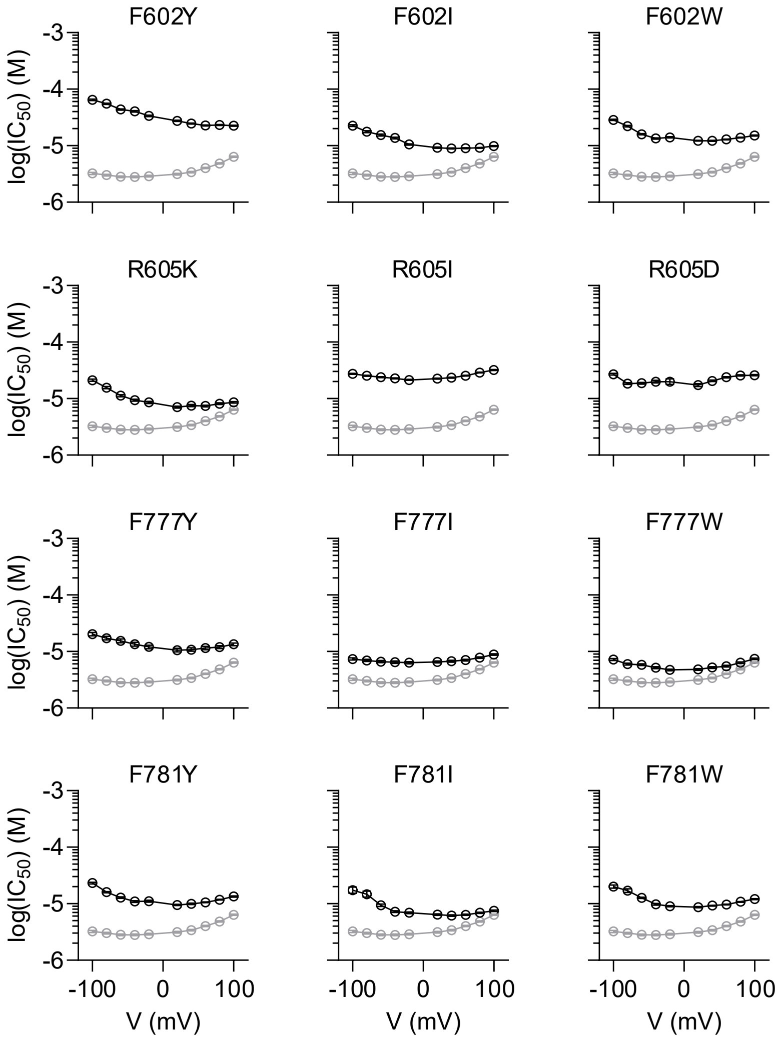


**Supplementary Fig. 14. $IC_{50}$ values of side chain mutagenesis on honokiol administration.** $IC_{50}$ values of TMEM16A whole-cell currents upon honokiol administration on the extracellular side. Intracellular $Ca^{2+}$ was fixed at 300 nM. $IC_{50}$ values were calculated by fitting to the Hill equation. Data are expressed as the estimated value ± 95% confidence interval. Source data are provided as Source Data file.

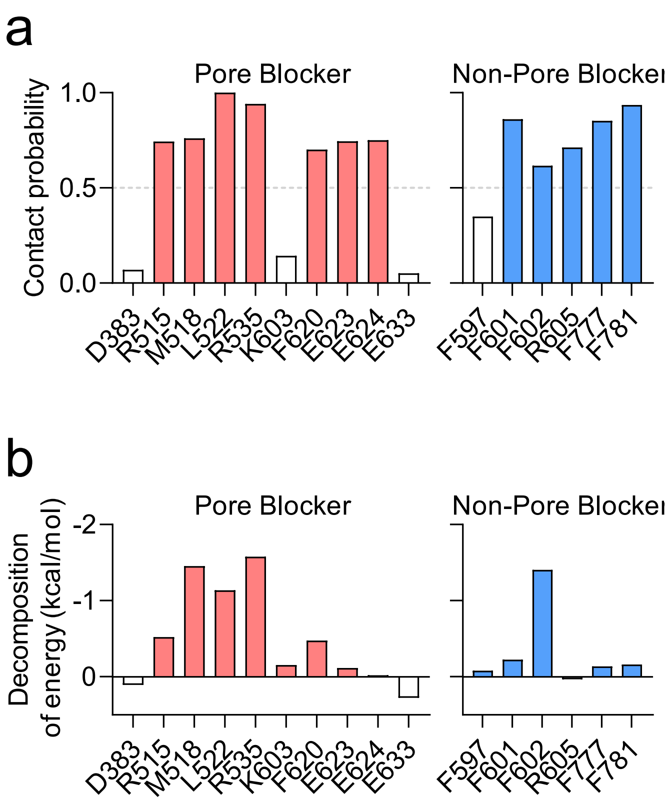


**Supplementary Fig. 15. Contact probability and decomposition of energy of magnolol and honokiol in the pore and non-pore pockets.**

**a.** Contact probability of pore and non-pore blockers. The contact probability was calculated when the distance between magnolol and honokiol to the residue was <4 Å. Residues with >0.5 contact probabilities are colored in red or blue. **b.** Decomposition of energy calculated by MM/GBSA (Molecular Mechanics with Generalized Born and Surface Area Solvation)[58] in MD simulation trajectories. Residues with negative decomposition of energy are colored in red or blue. Data are expressed as the mean. Source data are provided as Source Data file.

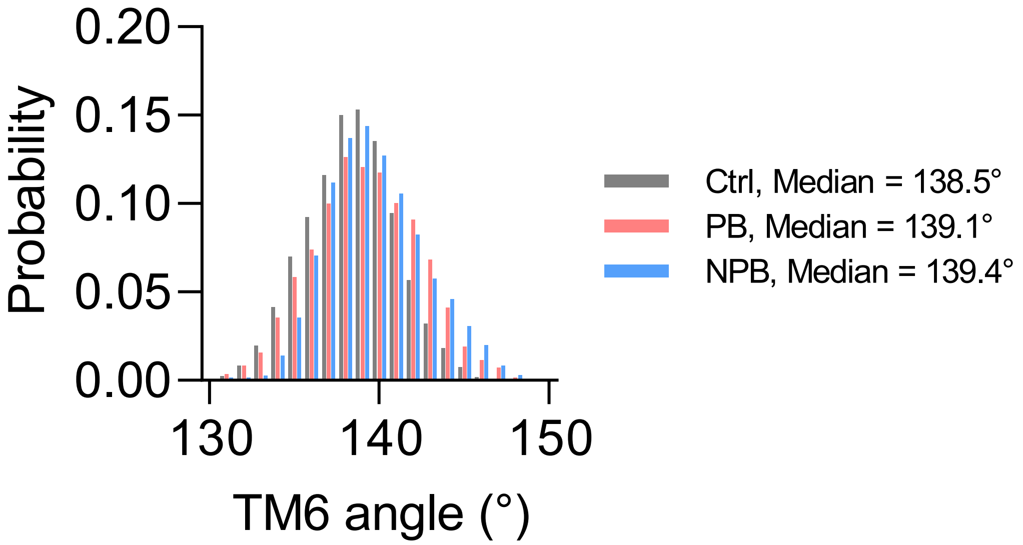


**Supplementary Fig. 16. TM6 angle in drug-free control state, pore blocker (magnolol)-bound state, and non- pore blocker (honokiol)-bound state.**

Angle of TM6 in MD simulation trajectories. The TM6 angle was calculated using VMD[55]. PB and NPB represent pore blocker-bound TMEM16A and non-pore blocker-bound TMEM16A, respectively. Source data were provided as source data files.

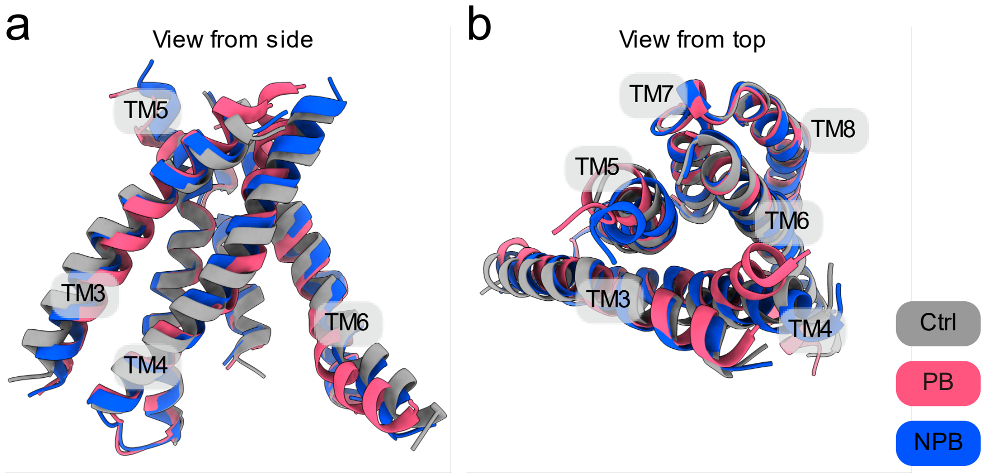


**Supplementary Fig. 17. Rearrangement of pore-lining TM domains in the presence of magnolol (pore blocker) and honokiol (non-pore blocker).**

**a.** Overlay structure from the MD simulations. Only pore-forming domains (TM3-8) are shown for clarity. View from side. **b.** View from top. PB and NPB represent pore blocker-bound TMEM16A and non-pore blocker-bound TMEM16A, respectively.

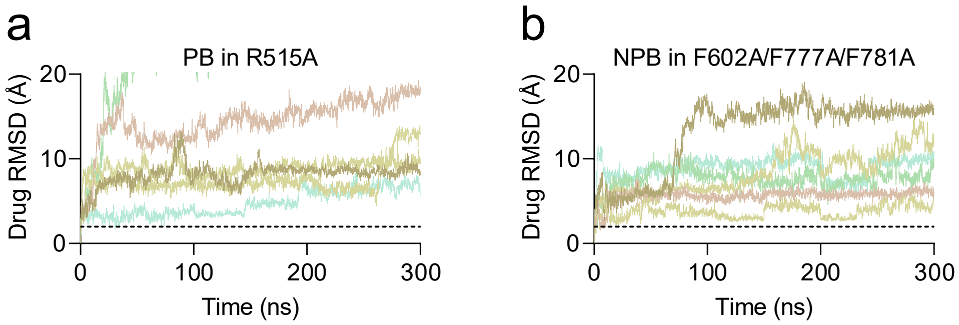


**Supplementary Fig. 18. Unstable binding of the pore and non-pore blockers in R515A and F602A/F777A/F781A mutants.**

**a.** RMSD calculation of pore blocker (PB; magnolol) to R515A mutant in MD simulations. b. RMSD calculation of non-pore blocker (NPB; honokiol) to F602A/F777A/F781A triple mutant in MD simulations. In **a** and **b**, RMSD of drugs were maintained around 2 Å in the WT MD simulations (Fig. 5b), which is indicated as dotted black line.

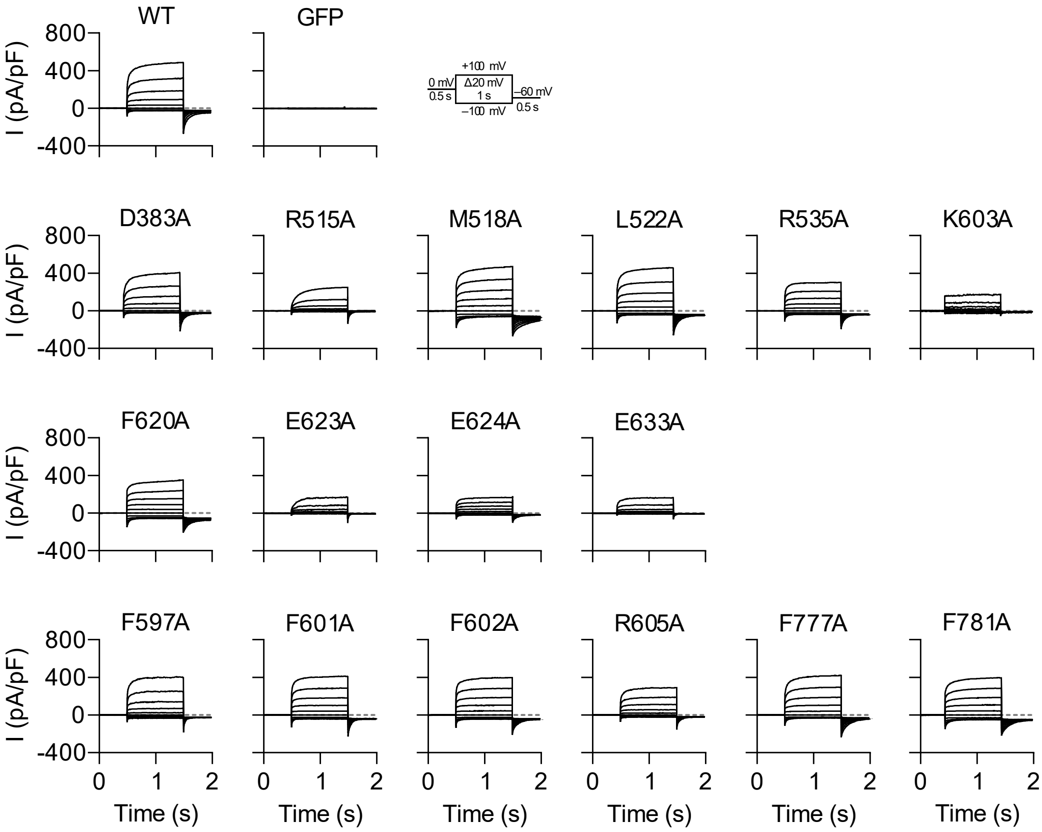


**Supplementary Fig. 19. Whole-cell current traces of the wild type and mutant TMEM16A.** Whole-cell currents of WT, pore-pocket mutants, and non-pore pocket mutants. Currents were evoked by 0 mV pre-pulse for 0.5 s, –100 to 100 mV test pulses with 20 mV increment for 1 s, and –60 mV post-pulse for 0.5 s. The first row shows the WT and green fluorescent protein (GFP), the second and third rows show the pore pocket mutants, and the last row shows the non-pore pocket mutants. Source data are provided as Source Data file.

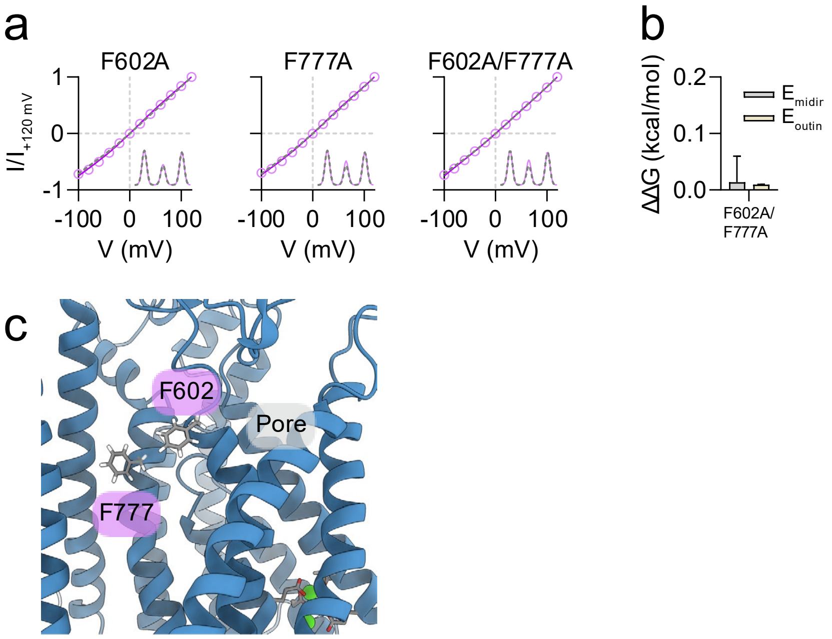


**Supplementary Fig. 20. Double-mutant cycle analysis for F602 and F777.**

**a.** Normalized current trace for single or double mutants of non-pore pocket residues. The pink circle represent the normalized current of each mutant, and the magenta line, the fit of the ion permeation model. The dotted gray line is the result of the WT. Energy barrier illustration is shown in inset. $n$ = 15 in each mutant. **b.** ΔΔG values calculated from double-mutant cycle analysis (see Supplementary Fig. 27 and Methods). The $E_{midin}$ value for F602A/F777A was 0.0139 ± 0.0833 kcal/mol and the $E_{outin}$ value was 0.0096 ± 0.0005 kcal/mol. **c.** Location of F602 and F777 in the outer pore region. Data are expressed as mean ± SEM in **a** and estimated value ± 95% confidence interval in **b**. Source data are provided as Source Data file.

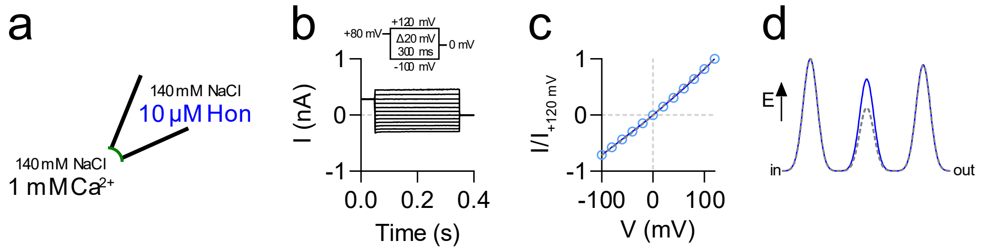


**Supplementary Fig. 21. Ion permeation model calculation with the non-pore blocker.**

**a.** Illustration of the inside-out patch-clamp for ion permeation model calculation with 10 μM honokiol. 1 mM $Ca^{2+}$ was used to activate TMEM16A in symmetrical 140 mM NaCl solutions. **b.** Current trace of WT TMEM16A activation curve in the presence of honokiol. Currents were evoked by 80 mV pre-pulse for 50 ms, followed by –100 to 120 mV test pulses for 300 ms. **c.** Normalized current trace. The blue circle represents the normalized current, and the blue line is the fit to the ion permeation model (see Methods). The dotted gray line represents the result of the drug-free state. *n* = 8. **d.** Energy barrier of WT TMEM16A in the presence of a non-pore blocker. Energy barriers were calculated by fitting the results from **c** to the ion-permeation model. $\Delta E_{midin}$ is –0.3611 ± 0.0002 kcal/mol and $\Delta E_{outin}$ is –0.1060 ± 0.0000 kcal/mol. The dotted gray line represents the result of the drug-free state. Source data are provided as Source Data file.

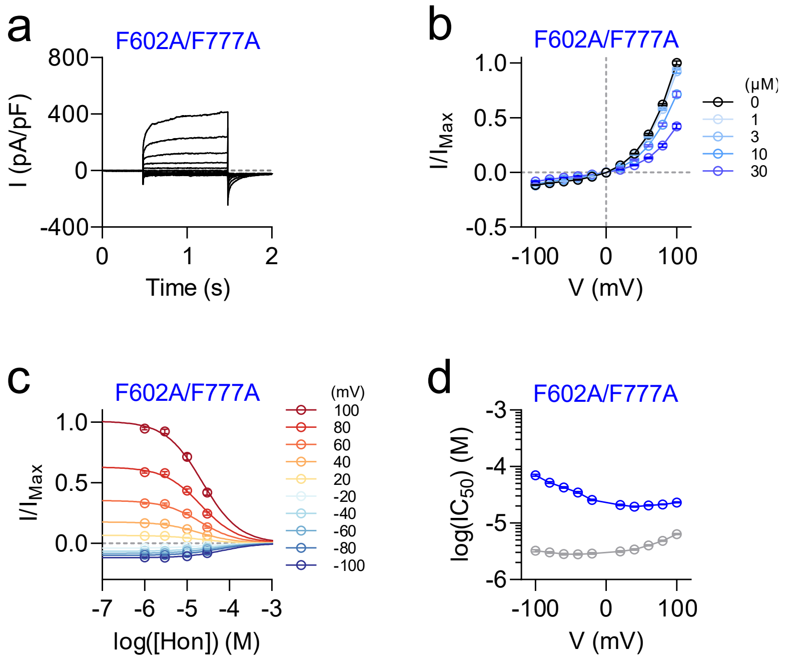


**Supplementary Fig. 22. Gating property and honokiol inhibition of F602A/F777A mutant.** **a.** Whole-cell currents of WT, pore-pocket mutants, and non-pore pocket mutants. Currents were evoked by 0 mV pre-pulse for 0.5 s, –100 to 100 mV test pulses with 20 mV increment for 1 s, and –60 mV post-pulse for 0.5 s. Intracellular $Ca^{2+}$ was fixed at 300 nM. **b.** Current-voltage relationships of TMEM16A whole-cell currents upon honokiol administration on the extracellular side. **c.** Dose-response relationships of TMEM16A whole-cell currents upon honokiol administration on the extracellular side. Curves are fit to the Hill equation. **d**. $IC_{50}$ values of TMEM16A whole-cell currents upon honokiol administration on the extracellular side. $IC_{50}$ values were calculated by fitting to the Hill equation. Data are expressed as the mean ± SEM in **b**, **c** and estimated values ± 95% confidence interval in **d**. Source data are provided as Source Data file.

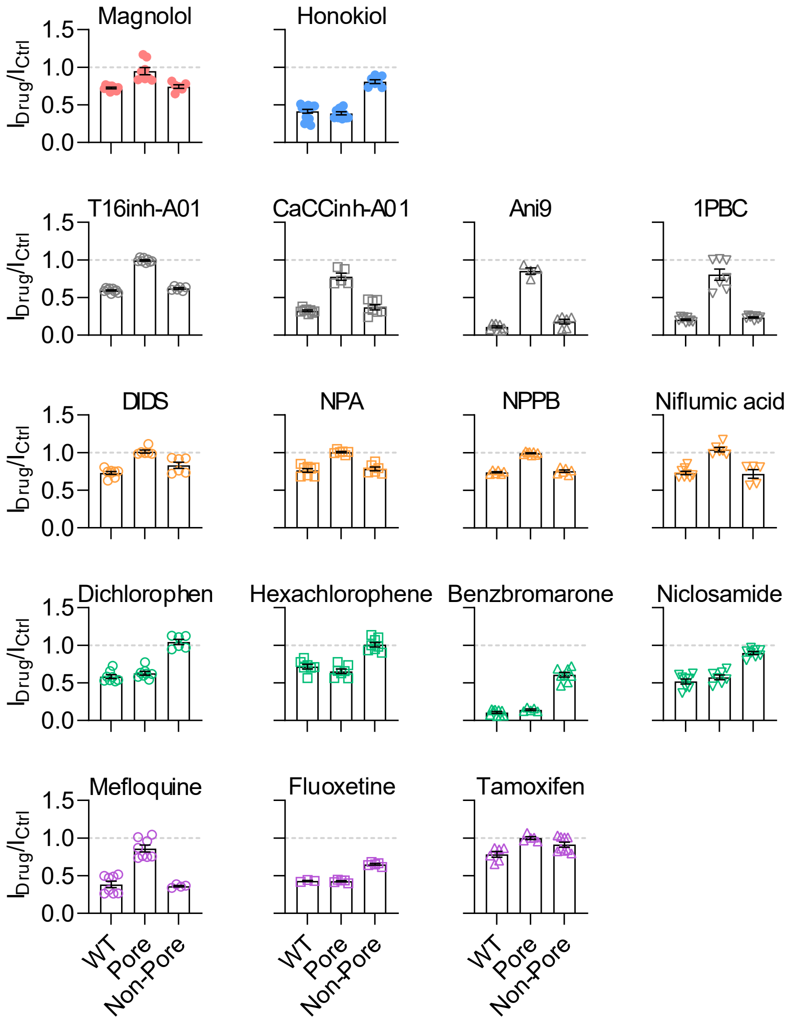


**Supplementary Fig. 23. Inhibition rate of the drugs on the WT, R515A (Pore) mutant, and F602A/F777A (Non-Pore) mutant.**

Each inhibitor was tested at 10 μM concentration on the extracellular side in the whole-cell patch-clamp configuration. Intracellular $Ca^{2+}$ concentration was fixed at 300 nM. The color codes for the pore and non-pore blockers are same as those in Fig. 8. Source data are provided as Source Data file.

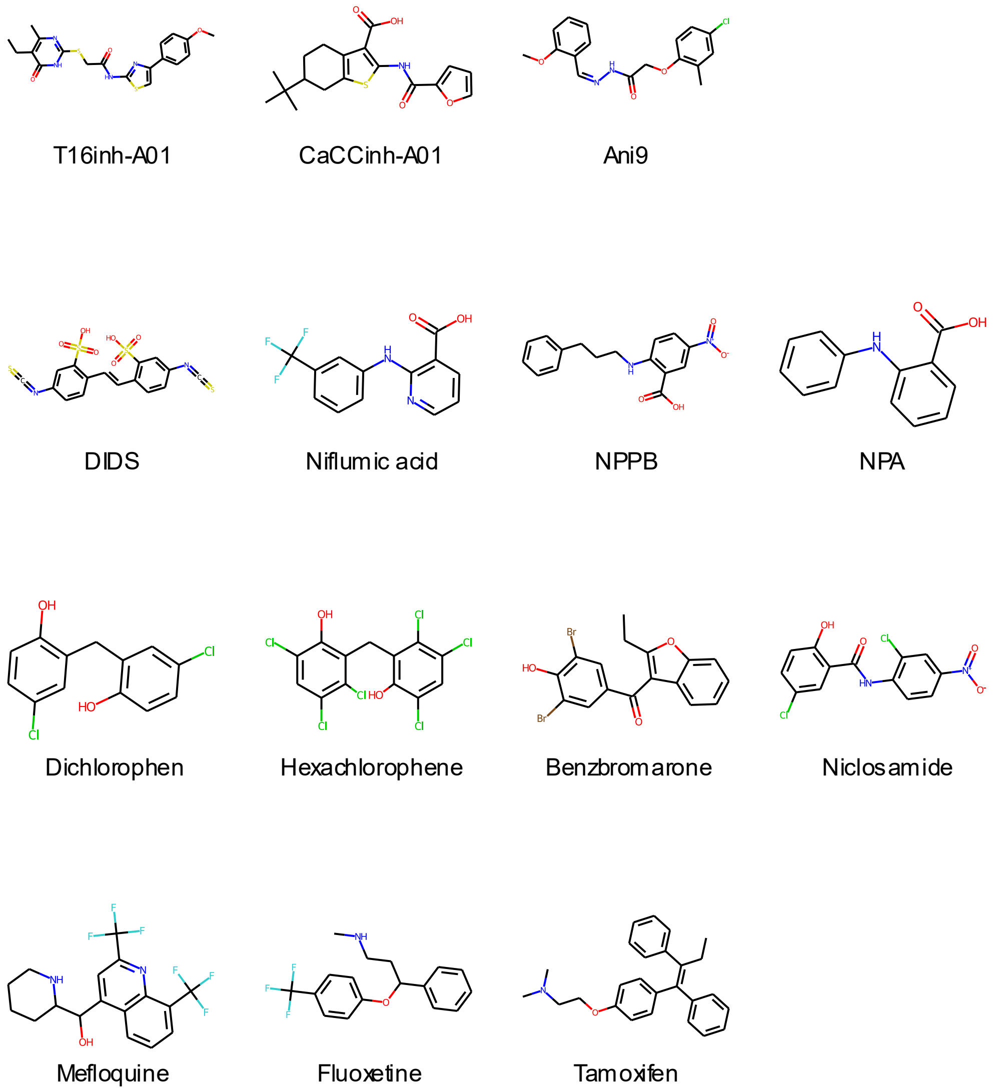


**Supplementary Fig. 24. 2D structures of TMEM16A inhibitors.**

2D structures of tested TMEM16A inhibitors in Fig. 8 and Supplementary Fig. 23.

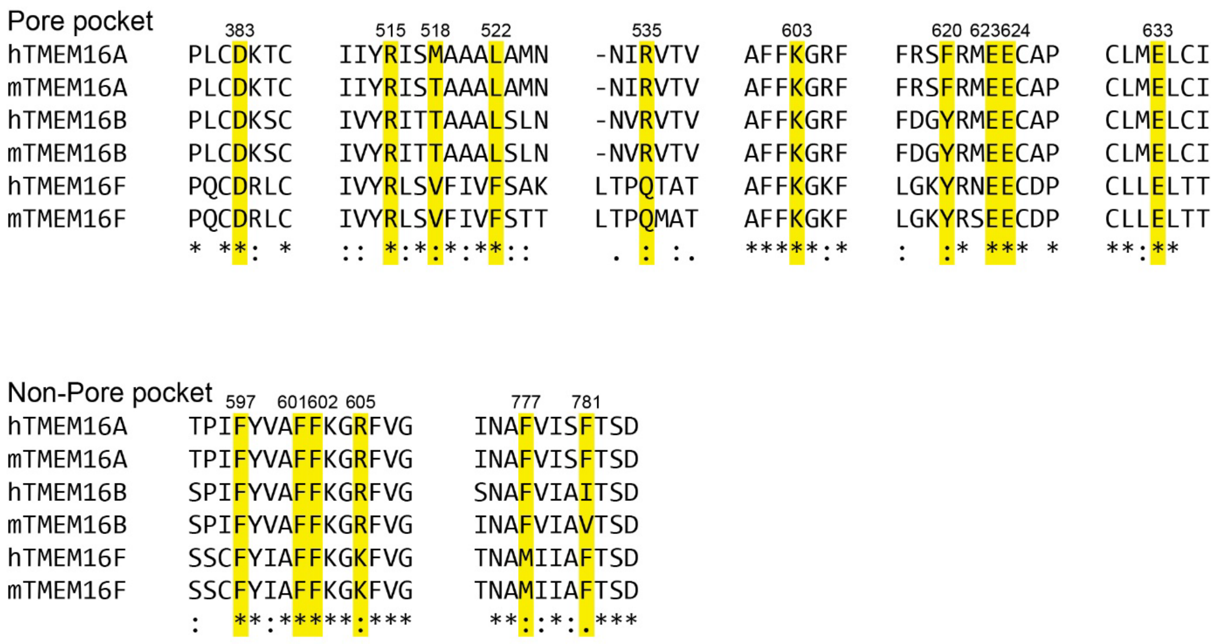


**Supplementary Fig. 25. Sequence alignment of TMEM16 pore and non-pore pocket regions.**

Sequence alignment of the pore and non-pore pocket residues in human and mouse TMEM16A, TMEM16B, and TMEM16F. Residue numbers denoted are reference to the human TMEM16A sequence.

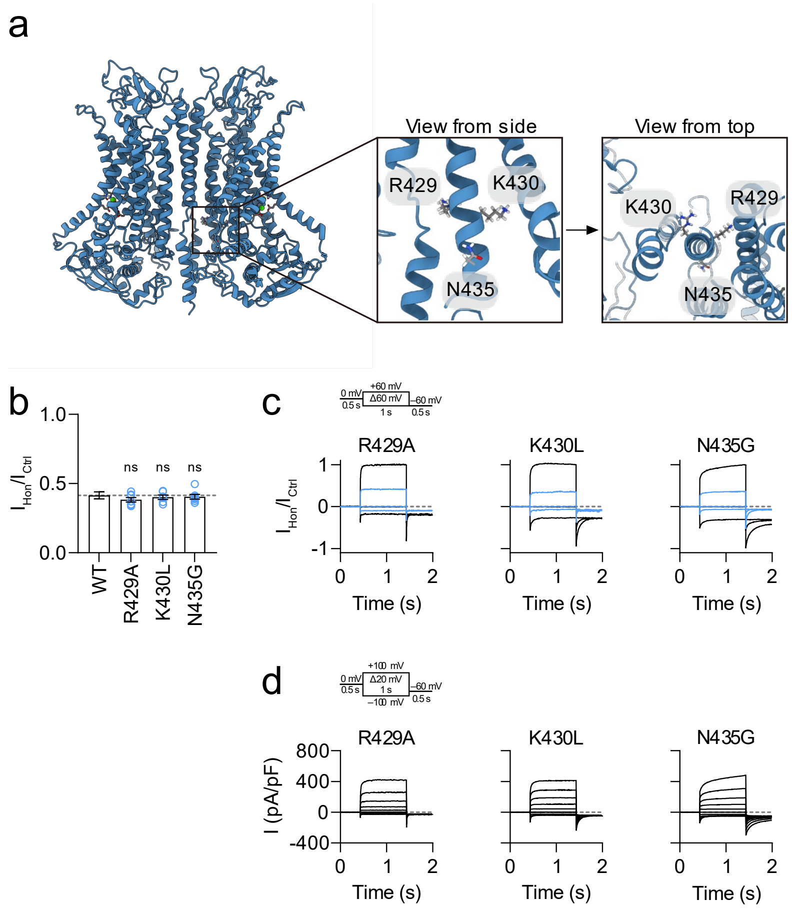


**Supplementary Fig. 26. Inhibition rate of honokiol on the binding site proposed by Wang *et al.*[41]**

**a.** Overall location of residues R429, K430, and N435. Note that these residues do not form a 3D pocket but rather stick out in different directions. **b.** Remaining currents for R429A, K430L, and N435G mutants. Intracellular $Ca^{2+}$ was fixed at 300 nM in a whole-cell patch-clamp configuration. Honokiol was applied to the extracellular side. $n$ = 15 for the WT and $n$ = 6–7 for each mutant. The gray line indicates the remaining normalized current for WT, which was 0.415. **c.** Inhibitory current traces for each mutant. Currents were evoked by 0 mV pre-pulse for 0.5 s, –60 or 0 or 60 mV test pulse for 1 s, and –60 mV post-pulse for 0.5 s. **d.** Whole-cell currents for each mutant. Currents were evoked by 0 mV pre-pulse for 0.5 s, –100 to 100 mV test pulses with 20 mV increment for 1 s, and –60 mV post-pulse for 0.5 s. Data are expressed as mean ± SEM. Source data are provided as Source Data file. $P$-values were calculated using one-way ANOVA, followed by Dunnett's post-hoc tests; ns, not significant.

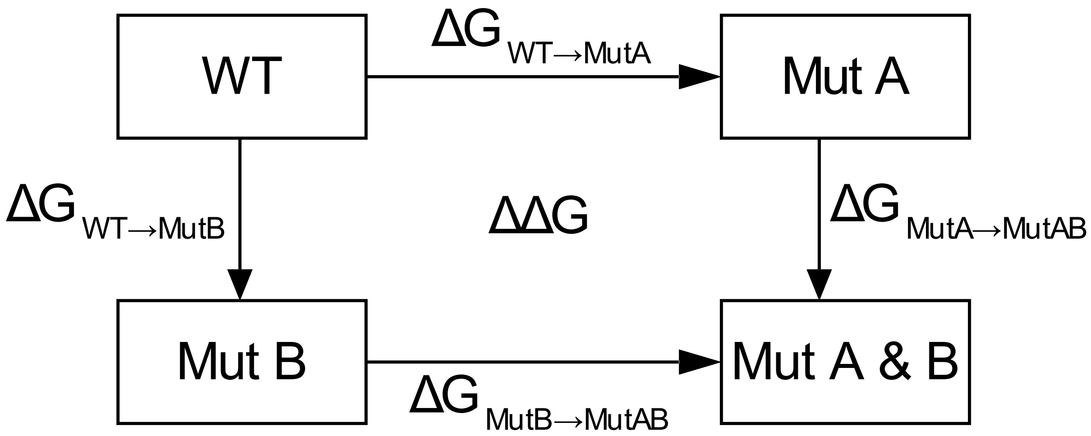


**Supplementary Fig. 27. Schematic diagram of double-mutant cycle analysis.**

Schematic diagram of double-mutant cycle analysis. WT is the wild-type protein, Mut A and Mut B are single mutants, and Mut A&B is the corresponding double mutant. ΔG is the free energy difference between two states, ΔΔG is the perturbation of the free energy between Mut A and Mut B.

**Supplementary Table 1. Clustering of docking poses by *k*-means algorithm**

| | Cluster | # of poses | Cluster type | % in 25 Å$^3$ | Note |
|---|---|---|---|---|---|
| Magnolol | 1 | 311 | Major | 0.99 | Pore pocket |
| | 2 | 281 | Major | 0.98 | Non-pore pocket |
| | 3 | 111 | Minor | | |
| | 4 | 93 | Minor | | |
| | 5 | 83 | Minor | | |
| | 6 | 70 | Minor | | |
| | 7 | 51 | Minor | | |
| Honokiol | 1 | 277 | Major | 1.00 | Non-pore pocket |
| | 2 | 258 | Major | 0.97 | Pore pocket |
| | 3 | 123 | Minor | | |
| | 4 | 103 | Minor | | |
| | 5 | 100 | Minor | | |
| | 6 | 77 | Minor | | |
| | 7 | 62 | Minor | | |

**Supplementary Table 2. List of primers used for mutagenesis**

| | |
|---|---|
| D383A | F: 5'- CCATGTGCCCGCTTTGCGCCAAGACCTGCAGCTACTG -3’<br>R: 5’- CAGTAGCTGCAGGTCTTGGCGCAAAGCGGGCACATGG -3' |
| R429A | F: 5'- CTTCATGGAGCACTGGAAGGCGAAACAGATGCGACTCAAC -3’<br>R: 5’- GTTGAGTCGCATCTGTTTCGCCTTCCAGTGCTCCATGAAG -3' |
| K430L | F: 5'- CATGGAGCACTGGAAGCGGCTACAGATGCGACTCAACTAC -3’<br>R: 5’- GTAGTTGAGTCGCATCTGTAGCCGCTTCCAGTGCTCCATG -3' |
| N435G | F: 5'- CGGAAACAGATGCGACTCGGCTACCGCTGGGACCTCAC -3’<br>R: 5’- GTGAGGTCCCAGCGGTAGCCGAGTCGCATCTGTTTCCG -3' |
| R515A | F: 5'- CTCGGCGTCATCATCTACGCAATCTCCATGGCCGCCGC -3’<br>R: 5’- GCGGCGGCCATGGAGATTGCGTAGATGATGACGCCGAG -3' |
| R515K | F: 5’- CTCGGCGTCATCATCTACAAAATCTCCATGGCCGCCGCC -3’<br>R: 5’- GGCGGCGGCCATGGAGATTTTGTAGATGATGACGCCGAG -3’ |
| R515I | F: 5’- CTCGGCGTCATCATCTACATAATCTCCATGGCCGCCGCC -3’<br>R: 5’- GGCGGCGGCCATGGAGATTATGTAGATGATGACGCCGAG -3’ |
| R515D | F: 5’- CTCGGCGTCATCATCTACGACATCTCCATGGCCGCCGCC -3’<br>R: 5’- GGCGGCGGCCATGGAGATGTCGTAGATGATGACGCCGAG -3’ |
| M518A | F: 5'- CATCATCTACAGAATCTCCGCGGCCGCCGCCTTGGCCATG -3'<br>R: 5'- CATGGCCAAGGCGGCGGCCGCGGAGATTCTGTAGATGATG -3' |
| M518T | F: 5’- ATCATCTACAGAATCTCCACGGCCGCCGCC -3’<br>R: 5’- GGCGGCGGCCGTGGAGATTCTGTAGATGAT-3’ |
| M518V | F: 5’- TCTACAGAATCTCCGTGGCCGCCGCCTTG -3’<br>R: 5’- CAAGGCGGCGGCCACGGAGATTCTGTAGA -3’ |
| M518I | F: 5’- CATCATCTACAGAATCTCCATAGCCGCCGCCTT -3’<br>R: 5’- AAGGCGGCGGCTATGGAGATTCTGTAGATGATG -3’ |
| L522A | F: 5'- CTCCATGGCCGCCGCCGCGGCCATGAACTCCTCC -3’<br>R: 5’- GGAGGAGTTCATGGCCGCGGCGGCGGCCATGGAG -3' |
| R535A | F: 5'- CGTGCGGTCCAACATCGCGGTCACAGTCACAGCC -3’<br>R: 5’- GGCTGTGACTGTGACCGCGATGTTGGACCGCACG -3' |
| F597A | F: 5'- CTACACCCCCATCGCTTACGTGGCGTTC -3’<br>R: 5’- GAACGCCACGTAAGCGATGGGGGTGTAG -3' |
| F601A | F: 5'- CCATCTTTTACGTGGCGGCCTTCAAAGGCCGGTTTG -3’<br>R: 5’- CAAACCGGCCTTTGAAGGCCGCCACGTAAAAGATGG -3' |
| F602A | F: 5’- CTTTTACGTGGCGTTCGCCAAAGGCCGGTTTGTTGG -3’<br>R: 5’- CCAACAAACCGGCCTTTGGCGAACGCCACGTAAAAG -3’ |
| F602Y | F: 5’- CATCTTTTACGTGGCGTTCTACAAAGGCCGGTTTGTTGGAC -3’<br>R: 5’- GTCCAACAAACCGGCCTTTGTAGAACGCCACGTAAAAGATG -3’ |
| F602I | F: 5’- CATCTTTTACGTGGCGTTCATCAAAGGCCGGTTTGTTGG -3’<br>R: 5’- CCAACAAACCGGCCTTTGATGAACGCCACGTAAAAGATG -3’ |
| F602W | F: 5’- CTTTTACGTGGCGTTCTGGAAAGGCCGGTTTGTTGG -3’<br>R: 5’- CCAACAAACCGGCCTTTCCAGAACGCCACGTAAAAG -3’ |
| K603A | F: 5'- CTTTTACGTGGCGTTCTTCGCAGGCCGGTTTGTTGGACGC -3’<br>R: 5’- GCGTCCAACAAACCGGCCTGCGAAGAACGCCACGTAAAAG -3' |

| | |
|---|---|
| K603R | F: 5’- CTTTTACGTGGCGTTCTTCAGAGGCCGGTTTGTTGGACGCC -3’<br>R: 5’- GGCGTCCAACAAACCGGCCTCTGAAGAACGCCACGTAAAAG -3’ |
| K603I | F: 5’- CTTTTACGTGGCGTTCTTCATAGGCCGGTTTGTTGGACGCC -3’<br>R: 5’- GGCGTCCAACAAACCGGCCTATGAAGAACGCCACGTAAAAG -3’ |
| K603D | F: 5’- CTTTTACGTGGCGTTCTTCGACGGCCGGTTTGTTGGACGCC -3’<br>R: 5’- GGCGTCCAACAAACCGGCCTCTGAAGAACGCCACGTAAAAG -3’ |
| R605A | F: 5'- GTGGCGTTCTTCAAAGGCGCGTTTGTTGGACGCCCGGG -3’<br>R: 5’- CCCGGGCGTCCAACAAACGCGCCTTTGAAGAACGCCAC -3' |
| R605K | F: 5’- CGTGGCGTTCTTCAAAGGCAAGTTTGTTGGACGCCCGGGC -3’<br>R: 5’- GCCCGGGCGTCCAACAAACTTGCCTTTGAAGAACGCCACG -3’ |
| R605I | F: 5’- TGGCGTTCTTCAAAGGCATCTTTGTTGGACGCCCGGG -3’<br>R: 5’- CCCGGGCGTCCAACAAAGATGCCTTTGAAGAACGCCA -3’ |
| R605D | F: 5’- GTGGCGTTCTTCAAAGGCGACTTTGTTGGACGCCCGGGC -3’<br>R: 5’- GCCCGGGCGTCCAACAAAGTCGCCTTTGAAGAACGCCAC -3’ |
| F620A | F: 5'- GTGTACATTTTCCGTTCCGCCCGAATGGAAGAGTGTGC -3’<br>R: 5’- GCACACTCTTCCATTCGGGCGGAACGGAAAATGTACAC -3' |
| E623A | F: 5'- CCGTTCCTTCCGAATGGCAGAGTGTGCGCCAGGGG -3’<br>R: 5’- CCCCTGGCGCACACTCTGCCATTCGGAAGGAACGG -3' |
| E623D | F: 5’- CCGTTCCTTCCGAATGGACGAGTGTGCGCCAGGGGGC -3’<br>R: 5’- GCCCCCTGGCGCACACTCGTCCATTCGGAAGGAACGG -3’ |
| E623I | F: 5’- CCGTTCCTTCCGAATGATAGAGTGTGCGCCAGGG -3’<br>R: 5’- CCCTGGCGCACACTCTATCATTCGGAAGGAACGG -3’ |
| E623K | F: 5’- CCGTTCCTTCCGAATGAAAGAGTGTGCGCCAGG -3’<br>R: 5’- CCTGGCGCACACTCTTTCATTCGGAAGGAACGG -3’ |
| E624A | F: 5'- GTTCCTTCCGAATGGAAGCGTGTGCGCCAGGGGGCTG -3’<br>R: 5’- CAGCCCCCTGGCGCACACGCTTCCATTCGGAAGGAAC -3' |
| E633A | F: 5'- CAGGGGGCTGCCTGATGGCGCTATGCATCCAGCTCAG -3’<br>R: 5’- CTGAGCTGGATGCATAGCGCCATCAGGCAGCCCCCTG -3' |
| F777A | F: 5'- GCTGTCATCATCAATGCCGCCGTGATCTCCTTCACGTC -3’<br>R: 5’- GACGTGAAGGAGATCACGGCGGCATTGATGATGACAGC -3' |
| F777Y | F: 5’- CTGTCATCATCAATGCCTACGTGATCTCCTTCACGTC -3’<br>R: 5’- GACGTGAAGGAGATCACGTAGGCATTGATGATGACAG -3’ |
| F777I | F: 5’- CTGTCATCATCAATGCCATCGTGATCTCCTTCACG -3’<br>R: 5’- CGTGAAGGAGATCACGATGGCATTGATGATGACAG -3’ |
| F777W | F: 5’- GCTGTCATCATCAATGCCTGGGTGATCTCCTTCACGTCTG -3’<br>R: 5’- CAGACGTGAAGGAGATCACCCAGGCATTGATGATGACAGC -3’ |
| F781A | F: 5'- CAATGCCTTCGTGATCTCCGCCACGTCTGACTTCATCCCG -3’<br>R: 5’- CGGGATGAAGTCAGACGTGGCGGAGATCACGAAGGCATTG -3' |
| F781Y | F: 5’- CAATGCCTTCGTGATCTCCTACACGTCTGACTTCATCCCGC -3’<br>R: 5’- GCGGGATGAAGTCAGACGTGTAGGAGATCACGAAGGCATTG -3’ |
| F781I | F: 5’- CAATGCCTTCGTGATCTCCATCACGTCTGACTTCATCCC -3’<br>R: 5’- GGGATGAAGTCAGACGTGATGGAGATCACGAAGGCATTG -3’ |
| F781W | F: 5’- GCCTTCGTGATCTCCTGGACGTCTGACTTCATCC -3’<br>R: 5’- GGATGAAGTCAGACGTCCAGGAGATCACGAAGGC -3’ |
